# A conformational-dependent interdomain redox relay at the core of Protein Disulfide Isomerase activity

**DOI:** 10.1101/2023.02.24.529848

**Authors:** Eduardo P. Melo, Soukaina El-Guendouz, Cátia Correia, Fernando Teodoro, Carlos Lopes

## Abstract

Protein disulfide isomerases (PDIs) are a family of molecular chaperones resident in the endoplasmic reticulum (ER) emerging as important factors in disease. In addition to an holdase function, some members catalyse disulfide bond formation and isomerization, a crucial step for native folding and prevention of aggregation of misfolded proteins. PDIs are characterized by a modular arrangement of thioredoxin-like domains, with the canonical, first identified PDIA1, organized as four thioredoxin-like domains forming a horseshoe with two active sites at the extremities. Using two fluorescent redox sensors, roGFP2 and HyPer, as client substrates either unfolded or native, and the *in vitro* reconstitution of the full pathways of oxidative protein in the ER, we clarified important aspects underlying the catalytic cycle of PDIA1. The N-terminal *a* active site is the main oxidant of thiols and can transfer electrons to the C-terminal *a’* active site relying on the redox-dependent conformational flexibility of PDIA1 that allows the formation of an interdomain disulfide bond. The *a’* active site act then as a crossing point to redirect electrons to the ER downstream oxidases or back to client proteins. The two active sites of PDIA1 work cooperatively as an interdomain redox relay that explains PDIA1 oxidative activity to form native disulfides and PDIA1 reductase activity to resolve scrambled disulfides. Moreover, this mechanism reveals a new rational for shutting down oxidative protein folding under ER redox imbalance or when the levels of unfolded proteins and folding intermediates exceed the folding capacity of the system.

## Introduction

The endoplasmic reticulum (ER) is responsible for the manufacturing, folding and assembling of about one-third of the cell protein repertoire (Ghaemmaghami et al., 2003). A key post-translational modification for many of these proteins is disulphide bond formation, the so-called oxidative protein folding, which stabilize proteins and permit redox control of activity in some cases. Spontaneous thiol-disulphide exchange is kinetically not competent on the folding timescale and therefore disulphide bond formation must be accelerated by catalytic machineries (Nagy, 2013). Although direct chemical catalysis by low molecular weight electron carriers such as hydrogen peroxide, dehydroascorbate and vitamin K may contribute to low extent for disulphide bond formation (Ruddock, 2012; Margittai et al., 2009; Hatahet and Ruddock, 2009), kinetic competence and accuracy is mostly assured by enzymatic mechanisms. These enzymatic machineries should even be more relevant in oxidizing compartments such as the ER where scrambled disulphides can be formed and should be reduced or isomerized to form native disulphides and prevent misfolding and aggregation. Some enzymes have been identified as being involved in oxidative protein folding in the ER. Quiescin-sulfhydryl oxidase, particularly the ER-localized fraction, could play a role in disulphide bond formation (Kodali and Thorpe, 2010) as well as glutathione peroxidases (Kanemura et al., 2020a; Nguyen et al., 2011) and vitamin K epoxide reductase (Schulman et al., 2010). However, it seems relatively consensual that the two most important enzymatic pathways are the PDI-ERO1α (protein disulphide isomerase-endoplasmic reticulum oxidase 1) and the PDI-PRDX4 (PDI-peroxiredoxin IV) pathways (Zito et al., 2010; Tavender et al., 2010). For both, PDI acts as the first electron acceptor transferring the disulphide bond to the client protein. Electrons are then shuttled from PDI to ERO1α with molecular oxygen acting as final electron acceptor or alternatively to PRDX4 which reduces hydrogen peroxide to form water. Involvement of PDI in several pathological conditions gives further relevance to its mechanism of action. Altered expression of PDIs is a prominent and common feature of neurodegenerative diseases (Andreu et al., 2012; Woehlbier et al., 2016) including prion disease (Hetz et al., 2005). PDI mediates cleavage of disulfide bonds in a glycoprotein required for HIV-1 cellular invasion (Barbouche et al., 2003). PDI expression is upregulated in several cancers and is associated with clinical outcomes (Xu et al., 2013).

There are at least 21 PDI-family members in the ER containing one or more thioredoxin-like domains but only a few can catalyse thiol-disulphide exchange reaction (Andreu et al., 2012). Among them, the widely studied canonical PDIA1 (UniProt P07237) is abundant, particularly in secretory tissues such as liver, pancreas and placenta (Freedman et al., 2017). It can catalyse thiol-disulfide oxidation, reduction and isomerization, the last of which occurs directly through intramolecular disulfide rearrangement or through cycles of reduction and oxidation (Ellgaard and Ruddock, 2005). PDIA1 comprises two thioredoxin-like catalytic domains, *a* and *a’*, which are separated by two thioredoxin-like non-catalytic domains, *b* and *b’*, organized sequentially as *a-b-b’-x-a’* to form a horseshoe shape with *x* being a linker between *b’* and *a’* (Ellgaard and Ruddock, 2005; Hatahet and Ruddock, 2009). The catalytic domains contain a characteristic WCGHC active-site motif and display cooperativity (Fredman et al., 2017). The *b’* domain provides the primary peptide binding site but for larger substrates all domains contribute to the binding site (Klappa et al., 1998). The linker region *x* seems to provide flexibility between the two arms of the horseshoe shape of PDIA1 containing the catalytic domains. The *x* linker can assume at least two different conformations and should adopt alternative conformations during the functional cycle of PDIA1 action (Nguyen et al., 2008; Yang et al., 2014). There is a more compact PDIA1 conformer where a hydrophobic pocket in the *b’* domain is capped by the *x* linker inhibiting substrate binding, and an open conformer with the two arms containing the catalytic sites placed more distantly. Whether the more compact conformer is simply a mechanism to inhibit substrate binding or part of the catalytic cycle of PDIA1, representing a post-releasing substrate step, requires further clarification. Also, the mechanistic details explaining the cooperativity between the two active sites of PDIA1 and their functional interplay with downstream oxidoreductases in the ER require further insight.

In this work, we have used two fluorescent redox sensors, roGFP2 and HyPer, as client proteins to obtain new mechanistic insight on the PDIA1 catalytic cycle and on its interplay with ERO1α and PRDX4. Monitoring two different processes in the fluorescent redox sensors, refolding and oxidation, and further structural insight of PDIA1 indicate disulfide exchange between the two active sites as the core mechanism of its activity and show the role of the *a’* active site as the crossing point either to resolve scrambled disulfide bonds or to transfer electrons to downstream oxidases.

## Material and Methods

### Proteins

Recombinant human PDIA1 (PDI A1 18–508, plasmid hPDI(18-508)pTrcHis-A), recombinant mouse PRDX4 (residues 37-274, plasmid pRSET_A_mPRDX4[37-274]), roGFP2 (roGFP2_pET-15b) and HyPer (HyPer_pSmt3_pET28b) were expressed in *E.coli* BL21 (DE3) strain upon IPTG (isopropyl B-D-1-thiogalactopyranoside) induction and purified as described in Correia et al., 2020. Mouse ERO1α (residues 23-464) poses solubility issues and therefore a fusion with glutathione S transferase (GST) through the Smt3 cleavage sequence was used (plasmid mERO1α[23-464]_pGS). *E. coli* Rosetta (DE3) cells were used to express the fusion ERO1α-GST and cells were disrupted by sonication after resuspension in PBS supplemented with a cocktail of protease inhibitors. The supernatant of the cell lysate was incubated overnight at 4°C on a wheel with Glutathione Sepharose 4B resin (GSTrap 4B, GE Healthcare). After several washes, the Ulp protease was added and cleavage of the Smt3 linker was done on sepharose beads. The GST-Smt3 linker remains on the beads and eluted ERO1α was load onto a Nickel column to remove the His-tag Ulp protease. The Chinese hamster binding immunoglobulin protein (BiP, residues 27-654, plasmid haBiP-27-654_pQE10) was expressed and purified as described elsewhere (Preissler et al., 2017).

### PDIA1 mutagenesis

PDIA1 WT backbone (hPDI(18-508)pTrcHis-A) and the quick change method were used to obtain PDIA1 mutants (see primers list in Table S1, supplementary material). Briefly, primers containing one or two mutagenized codons were used in a PCR with the following steps: 1 min at 95°C, 20 cycles comprising 1 min at 95°C, 1 min at 65 to 70°C (depending on the annealing temperature of primers) and 7 min at 72°C with a final step at 72°C for 10 min. PCR products were incubated with DpnI for 2 h at 37°C before transforming competent DH5α *E. coli* cells. PDIA1 mutants with no attacking Cys (C53S-C397S), no resolving Cys (C56S-C400S) and no active site Cys (C53S-C56S-C397S-C400S), were generated after double digestion with Eco52I and EcoRI and fragments containing the desired mutations were purified from the gel and ligated with T4 ligase for 1 h at RT before transformation of DH5α *E. coli* cells. All the mutants were confirmed by DNA sequencing.

### Protein reduction

Proteins were reduced by incubation with 50 mM dithiothreitol (DTT), 1 h at room temperature, and DTT was removed by gel filtration on PD Minitrap G-25 columns (GE Healthcare) after column equilibration with strictly degassed buffer.

### Thermodynamic stability of roGFP2 and HyPer

The thermodynamic stability of roGFP2 and HyPer were measured after unfolding using increasing concentrations of guanidinium hydrochloride (GdnHCl) in Tris-HCl 50 mM buffer, pH 7.4, 150 mM NaCl and fluorescence excitation spectra were recorded with a Fluoromax 4, Horiba Scientific, as described in Correia et al., 2020.

### Redox potential of HyPer

Midpoint redox potential (E_0_’) of HyPer was determined after incubation with different ratios of 10 mM glutathione couple (GSH/GSSG for reduced and oxidised glutathione, respectively) in strictly degassed buffer using the Nernst equation as described in Hanson et al., 2004, Lohman and Remington, 2008. Fraction of HyPer reduced after equilibrium with the glutathione couple was determined from the excitation fluorescence intensity ratio 488/405 considering fully reduced and fully oxidised for 10 mM GSH and 10 mM GSSG, respectively, and a E_0_’ of −240 mV for the glutathione couple (Bekendam et al., 2016).

### HyPer and roGFP2 refolding catalysed by PDIA1

Refolding of HyPer (0.5 μM) and roGFP2 (0.25 μM) was measured in degassed Tris-HCl, 50 mM, pH 7.4, 150 mM NaCl buffer after dilution of the denaturant GdnHCl to 0.1-0.2 M, through the recovery of the fluorescence excitation spectra in the visible range using an emission wavelength of 530 nm. A quartz cuvette with 1 cm path length under magnetic stirring in a Fluoromax 4, Horiba Scientific, was used to record the excitation spectra and PDIA1 (5 μM) was added reduced or oxidised immediately after dilution of the chemical denaturant.

### HyPer and roGFP2 refolding and oxidation catalysed by PDIA1 and downstream ERO1α and PRDX4 oxidoreductases

Refolding and oxidation of HyPer and roGFP2 were measured in Tris-HCl, 50 mM, pH 7.4, 150 mM NaCl buffer as described above using PDIA1 (5 μM) plus 2 μM of ERO1α or PRDX4 added immediately after PDIA1 or later after some degree of refolding catalysed by PDIA1 was accomplished. In the case of PRDX4, glucose (2.5 mM) and glucose oxidase (10 mU/mL) were also added to the reaction mixture as an exogenous source of hydrogen peroxide to act as final electron acceptor for PRDX4.

### HyPer and roGFP2 oxidation catalysed by PDIA1 and downstream ERO1α and PRDX4 oxidoreductases

Oxidation of native HyPer (0.5 μM) and native roGFP2 (0.25 μM) by PDIA1 was measured in Tris-HCl, 50 mM, pH 7.4, 150 mM NaCl buffer through the fluorescence intensity ratio 488/405 nm and 400/492 nm of the excitation spectra, respectively. PDIA1 (5 μM) was added in the reduced state to prevent substrate oxidation unless the downstream oxidase ERO1α or PRDX4 (2 μM in the oxidised state) was added to the reaction mixture. For PRDX4, glucose (2.5 mM) and glucose oxidase (10 mU/mL) were also added to the reaction mixture.

### SDS-Page

Acrylamide/Bisacrylamide 12.5 % SDS gels were ran under reducing (100 mM DTT) and non-reducing conditions (no DTT). Samples loaded in non-reducing conditions were first reacted with 20 mM N-Ethylmaleimide (NEM) for 15 min on ice to prevent further cysteine oxidation.

### SwitchSENSE PDIA1 biophysical analysis

PDIA1 was immobilized through the HisTag onto a Nickel loaded nitrilotriacetic acid (NTA) group attached to a 48 bp DNA strand complementary to a DNA strand-fluorescent labelled, immobilized on the surface of the biochip (Tris-NTA kit CK-TN-1-B48 and the biochip MPC-48-1-R1-S from dynamic BIOSENSORS). The upward and downward motion of the double-strand DNA molecule under applied alternating electric potentials can be followed through the quenching of fluorescence provided by the gold surface of the biochip. The motion of DNA occurs on a microsecond time scale and the attachment of a protein to the surface-distal end of the DNA slows down the switching dynamics providing a measure dependent on the hydrodynamic volume and conformation of the protein (Langer et al., 2013). A dynamic response (DR) parameter, which equates the area under the fluorescence curve when switching the DNA lever upward at high frequency (10 kHz), correlates with the velocity of switching. High DR values indicate fast switching because maximum fluorescence is attained faster when the DNA lever tilt away from the gold quenching surface of the biochip. Switch-sense biophysical analysis was done with 200 nM of PDIA1 in TE-40 buffer (10 mM tris, 40 mM NaCl, 0.05 mM EDTA, 0.05 mM EGTA and 0.05 % Tween 20).

### Structural characterization of PDIA1

Tryptophan fluorescence emission spectra of 1 μM PDIA1 were measured in 50 mM Tris-HCl, 150 mM NaCl, pH 7.4 buffer on a Fluoromax 4, Horiba Scientific, using a 1 cm pathlength quartz cuvette, excitation at 296 nm and emission from 310 to 450 nm, slits 5 nm, and three scans were averaged for each spectrum. Far-UV circular dichroism (CD) spectra were recorded on a Jasco J-815 CD spectrometer for 1 μM PDIA1 in water using a 1 mm path length quartz cuvette, 1 nm step, 2 nm slits and five scans were averaged for each spectrum.

## Results

### Refolding, stability and fluorescence properties of the redox sensors roGFP2 and HyPer

RoGFP2 is a derivative of the green fluorescent protein containing a disulfide bond on the surface of the ß-barrel structure (Hanson et al., 2004). As a redox sensor, the excitation fluorescence intensity ratio 400/492 nm increases with oxidation from dithiol to disulfide. The engineered disulfide bond increases the stability of the native state and changes the unfolding pathway leading to the accumulation of an intermediate state (Figure S1A). When roGFP2 in the reduced state (roGFP2_RED_ herein) is unfolded by denaturant, it can refold to the native state upon dilution of the denaturant to acquire the fluorescent emissive state (Figure 1A, red spectrum). The final state after refolding is reduced as shown by the ratio 400/492 nm of 0.21 compared to a value of 0.17 for the standard native reduced (for clarity spectrum not shown in Fig. 1A). On contrary, if roGFP2 oxidised (roGFP2_OX_ herein) is chemically unfolded, no significant fluorescence emission is recovered after dilution of the denaturant, showing that roGFP2_OX_ cannot refold due to the conformational constraint imposed by the disulfide bond (Figure 1A, magenta spectrum and figure S2A). A reducing agent such as DTT at mM concentration can reduce the cufflink allowing refolding to the native fully reduced state (Figure S2B).

**Figure 1.**
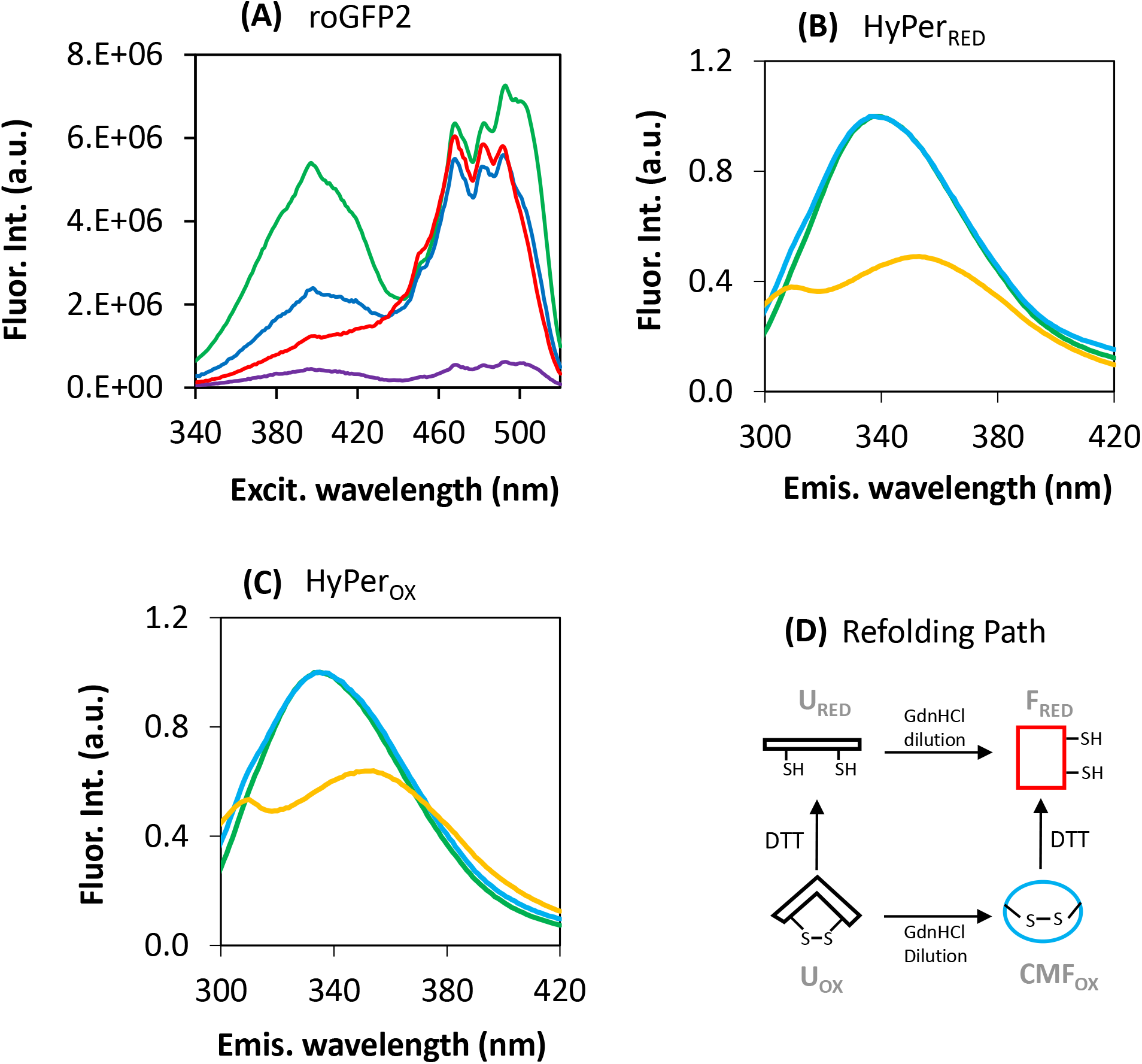
Panel (A) shows the fluorescence excitation spectra in the visible range of roGFP2. Chemically unfolded roGFP2_RED_ at 5.4M GdnHCl can refold and acquire visible fluorescence upon dilution of the denaturant to 0.2M ending up native and reduced (intensity ratio 400/492 nm of 0.21 for the red spectrum). On contrary, chemically unfolded roGFP2_OX_ cannot refold upon dilution of denaturant as shown by the low fluorescence recovered (magenta spectrum). Addition of PDIA1_RED_ (5μM) catalyzes the refolding of roGFP2_OX_ to the fluorescence emissive native structure and the final state is partly oxidized with a ratio 400/492 nm of 0.41 (blue spectrum). The ratio 400/492 nm for the oxidised native standard of roGFP2 is 0.71 (green spectrum) and for the reduced native standard is 0.17 (spectrum not shown for clarity). Panels (B) and (C) show the normalized tryptophan fluorescence emission spectra of HyPer in the UV range for reduced (B) and oxidised states (C). Green spectra are for native HyPer, light blue for HyPer chemically unfolded at 4.7M GdnHCl and refolded by dilution of denaturant to 0.1M and orange for HyPer unfolded in the preoxidation for the refoldingsence of GdnHCl. Panel (D): Schematic representation of the pathways for the refolding of HyPer and roGFP2 upon dilution of GdnHCl and addition of DTT. U stands for the unfolded state, F for the native state and CMF for the compact misfolded state (no fluorescence in the visible range but tryptophan burying) and subscripts RED and OX for reduced and oxidized, respectively.

HyPer is a circularly permutated variant of the yellow fluorescent protein engineered to contain the regulatory domain of the H_2_O_2_-sensing protein OxyR (Belousov et al., 2006). The introduction of this domain turns HyPer into a fluorescent ratiometric sensor that responds in a concentration dependent manner to H_2_O_2_ *in vitro* and in cells (Melo et al., 2017). The excitation fluorescence intensity ratio 488/405 nm of native HyPer increases with the formation of the disulfide bond on the OxyR domain. The contribution of this disulfide bond to HyPer stability was measured as 4.3 kcal/mol from the difference between the stability of HyPer oxidised (HyPer_OX_ herein) and Hyper reduced (HyPer_RED_ herein) (Figure S1B). As observed for roGFP2, chemically-unfolded HyPer can refold to the fully emissive native state when redox cysteines were reduced but cannot do it if the disulfide bond is formed (Correia et al., 2020). If the constraint imposed by the disulfide bond is removed by mM concentrations of DTT, then HyPer refolds to the native reduced emissive state. Interestingly, unfolded HyPer_OX_ cannot refold to the native state to acquire fluorescence in the visible range but collapses to a compact structure upon dilution of the denaturant. This conclusion is drawn from the fluorescence emission spectra of HyPer_OX_ in the UV range. HyPer unfolded emits in the UV range with a peak at 354 nm (Figure 1B and 1C, yellow spectra) as expected from tryptophan residues exposed to water (Lakowicz, 1999). Upon denaturant dilution, the peak for tryptophan emission occurs at 336 nm for both HyPer_OX_ and HyPer_RED_ matching the peak observed for native HyPer (Figure 1B and 1C, compare blue and green spectra). The emission peak revealed the same buried environment for tryptophan residues in HyPer_OX_ and HyPer_RED_ after refolding although fluorescence emission of the native state in the visible region can only be regained after refolding of HyPer_RED_. Protein hydrophobic collapse under native conditions is a well characterized phenomena in protein folding (Fersht, 2017). However, hydrophobic collapse channels HyPer_OX_ to a compact conformational trap from where it cannot refold to the native state. We defined this trap as a compact misfolded state (no fluorescence in the visible range but burying of tryptophan residues) which can mimetize protein conformations resulting from scrambled disulfide bonds. As well as the disulfide bond on unfolded HyPer and roGFP2, scrambled disulfide bonds should be resolved to avoid misfolding and prevent loss-of-function (Yang et al., 2015).

Summarizing, both HyPer_RED_ and roGFP2_RED_ refold to the native fluorescent emissive state upon dilution of denaturant (schematic representation in Figure 1D). With the redox-sensitive disulfide bond formed, neither HyPer_OX_ nor roGFP2_OX_ can refold to the native state because the disulfide bond acts like a cufflink and the conformational path to the native conformation is no longer accessible. However, dilution of denaturant promotes a protein hydrophobic collapse and channels the protein to a compact misfolded state with buried tryptophan residues.

### Refolding and disulfide bond formation on HyPer and roGFP2 by PDIA1

PDIA1, predominantly localized to the endoplasmic reticulum, acts as both chaperone and oxidoreductase (Cai et al., 1994; Song and Wang, 1995; Winter et al., 2002). Taking advantage of the fluorescent properties of HyPer and roGFP2, we have addressed the activity of PDIA1 on the formation/reduction of disulfide bonds and on the capability to promote protein refolding through its chaperone activity. PDIA1 in the reduced state (PDIA1_RED_ herein) but not in the oxidised state (PDIA1_OX_ herein) can catalyse the reduction of unfolded/misfolded roGFP2_OX_ and assist on protein folding (Figure 1A, blue spectrum, and figures S2C and S2D). The final state after refolding is native roGFP2 partly oxidised as shown by the ratio 400/492 nm of 0.41 compared to 0.71 and 0.17 for the fully oxidised and fully reduced standards, respectively (compare the blue and the green spectra in Figure 1A). As well as for roGFP2_OX_, refolding of HyPer_OX_ can be catalysed by PDIA1_RED_ but not by PDIA1_OX_ as shown previously (Correia et al., 2020). PDIA1_RED_ can resolve the compact misfolded state that accumulates when the oxidised protein is placed under native conditions. This capability to resolve kinetically trapped intermediates resulting from non-native disulfide bonds was reported for several proteins, including the human hormone chorionic gonadotropin and hen lysozyme (Huth et al., 1993; van den Berg et al., 1999; Kanemura et al., 2020b).

Refolding kinetics of HyPer_OX_ catalysed by PDIA1_RED_ increase with the concentration of PDIA1_RED_ (Figure 2A). Moreover, kinetics increased hyperbolically with the concentration of PDIA1_RED_ as expected from enzyme kinetics (Figure S3). The ER chaperone BiP (binding immunoglobulin protein also known as GRP78) with no reductase activity is unable to catalyse refolding of HyPer_OX_ confirming that reduction of the disulfide bond is the key step for refolding (Figure 2A). The midpoint redox potential of native HyPer was determined as −260± 5 mV using the Nernst equation and the glutathione redox couple (Figure S4). For roGFP2, values in the range of −272 to −280 mV were reported (Hanson et al., 2004; Björnberg et al., 2006). According to these redox potentials, PDIA1 with an apparent redox potential of −206 mV for both active sites as determined by mass spectrometry (Bekendam et al., 2016) should oxidise HyPer and roGFP2, instead of reducing them as observed in our refolding assays. Indeed, oxidation of native roGFPiE_RED_ from dithiol to disulfide (Avezov et al., 2013, 2015) as well as of native HyPer_RED_ (Konno et al., 2015; Melo et al., 2017) catalysed by PDIA1_OX_ was recorded. Native rodGFP2_RED_ can also be oxidised by PDIA1_OX_ (Figure S5). Capability of PDIA1 to reduce and refold HyPer_OX_ and roGFP2_OX_ (Figure 1A and 2A) indicates that the redox potential of the disulfide bond in the misfolded trap of the client proteins should be less negative than that of PDIA1. In nature, midpoint redox potentials of protein disulfides cover an extremely large range that spans the relevant physiological states. For unfolded proteins seems to be around −220 mV (Hatahet and Ruddock, 2009) with PDIA1 acting as an oxidant too. Compact misfolded states of HyPer_OX_ and roGFP2_OX_ accumulated after dilution of denaturant are surrogates of conformational states with scrambled disulfide bonds which should be more oxidising than those in native and unfolded proteins to be resolved by PDIA1.

**Figure 2.**
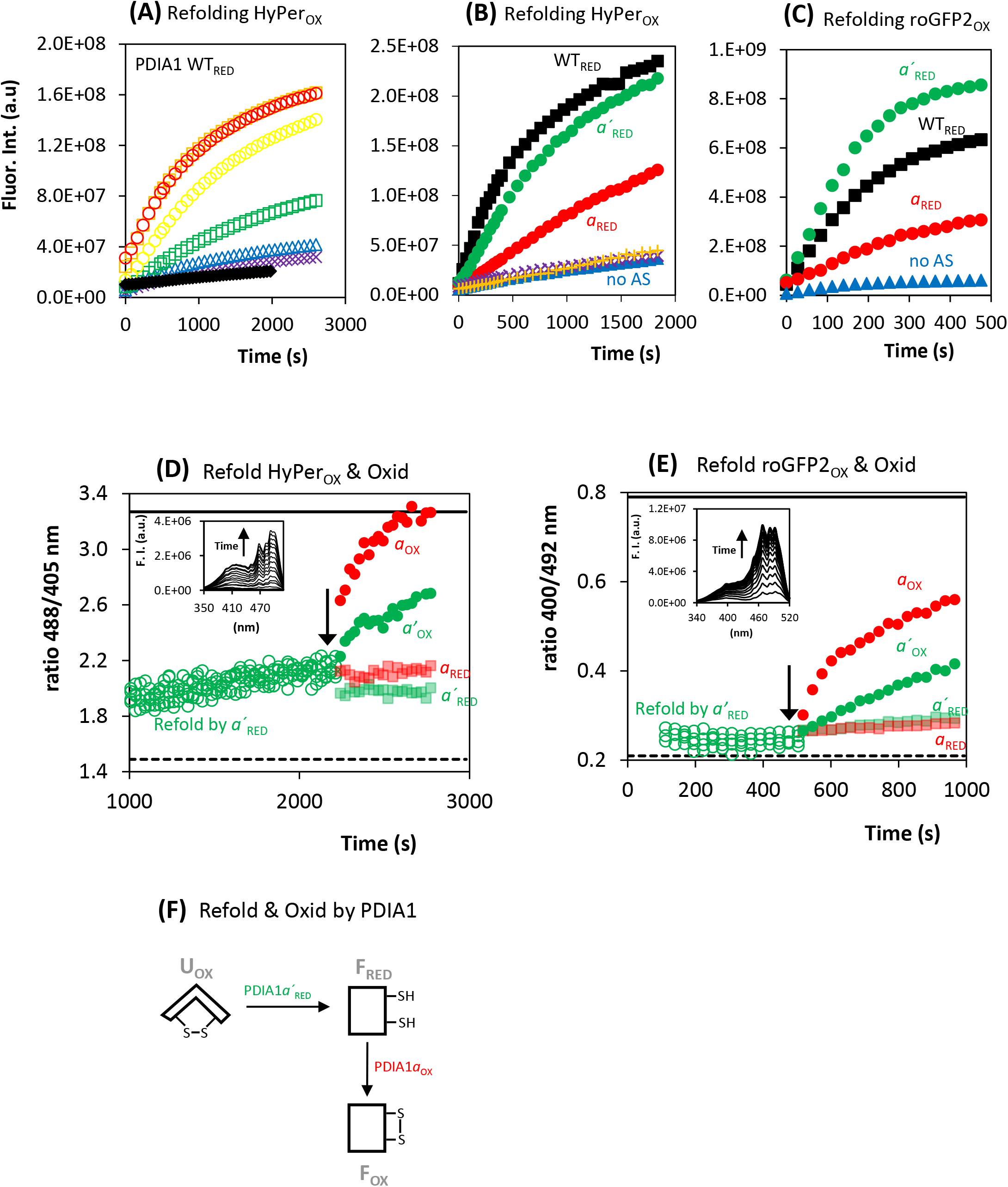
Panel (A) Refolding of HyPer_OX_ for different concentration of PDIA1_RED_ measured from the time-dependence of the integrated fluorescence intensity of each excitation spectra. No PDIA1_RED_ (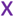), 0.5 (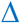), 1 (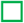), 2.5 (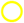), 5 (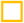) and 10 μM PDIA1_RED_ (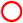). A control was run with the ER chaperone BiP (5 μM) plus 1 mM ATP (♦). Panel (B) Refolding of HyPer_OX_ by different mutants of PDIA1_RED_ (5 μM) measured from the time-dependence of the integrated fluorescence intensity of each excitation spectra. PDIA1 WT_RED_ (■), PDIA1*a*_RED_ (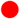), PDIA1*a’*_RED_ (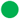), PDIA1 no active site cysteines (no AS) (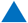), PDIA1 no attacking Cys (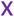) and no resolving Cys (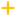). Panel (C) Refolding of roGFP2_OX_ measured from the time-dependence of the integrated fluorescence intensity of each excitation spectra. Refolding was catalyzed by PDIA1 WT_RED_ (■), PDIA1*a*_RED_ (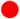), PDIA1*a’*_RED_ (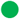) and PDIA1 no AS (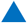). Initial rate constants of refolding were respectively 22.6×10^5^, 7.9×10^5^, 32.3×10^5^ and 2.5×10^5^ s^-1^. Panel (D) Excitation fluorescence intensity ratio (488/405 nm) reports dithiol oxidation for the refolding of HyPer_OX_ catalyzed by PDIA1*a*’_RED_ during the first 2200 s (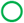) followed by the addition (pointed by arrow) of the PDIA1*a*_RED_ (fade 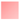), PDIA1*a*’_RED_ (fade 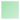), PDIA1*a*_OX_ (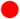) and PDIA1*a*’_OX_ (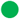). Dashed line at the bottom is the fluorescence ratio for the HyPer_RED_ standard and solid line at the top is the ratio for the HyPer_OX_ standard. Inset shows the excitation spectra of HyPer during refolding catalyzed by PDIA1*a’*_RED_ during the first 2200 s. Panel (E) Excitation fluorescence intensity ratio (400/492 nm) reports directly dithiol oxidation for the refolding of roGFP2_OX_ catalyzed by PDIA1*a*’_RED_ for the first 500 s (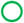) followed by the addition (pointed by arrow) of the PDIA1*a*_RED_ (fade 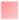), PDIA1*a’*_RED_ (fade 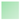), PDIA1*a*_OX_ (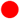) and PDIA1*a’*_OX_ (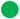). Dashed line at the bottom is the fluorescence ratio for the roGFP2_RED_ standard and solid line at the top is the ratio for the roGFP2_OX_ standard. Inset shows the excitation spectra of roGFP2 during refolding catalyzed by PDI*a’*_RED_ for the first 500 s. Panel (F): Schematic representation of the preferential pathway for refolding and oxidation of HyPer_OX_ and roGFP2_OX_ catalyzed by PDIA1*a*’_RED_ and PDIA1*a*_OX_, respectively. U stands for the unfolded state and F for the native state. Panels (B) to (E) show one representative assay from three independent experiments. PDIA1*a* (mutant C397S-C400S); PDIA1*a’* (mutant C53S-C56S); PDIA1 no AS (mutant C53S-C56S-C397S-C400S); PDIA1 no attacking cysteines (mutant C53S-C397S); PDIA1 no resolving cysteines (mutant C56S-C400S).

To address the role of each active site of PDIA1 on the refolding of HyPer_OX_ and roGFP2_OX_, we tested PDIA1 mutants with no cysteines in the *a* active site (C53S-C56S, herein named PDIA1*a’* mutant as it holds the *a’* active site), with no cysteines in the *a’* active site (C397S-C400S, named PDIA1*a*), with no attacking cysteines (C53S-C397S), with no resolving cysteines (C56S-C400S) and with none of the active site cysteines (C53S-C56S-C397S-C400S, named PDIA1 no active sites (no AS)). PDIA1*a*’_RED_ was almost as efficient as the WT_RED_ on the refolding of HyPer_OX_ and significantly more efficient than PDIA1*a*_RED_ (Figure 2B). Mutagenesis of attacking or resolving cysteines compromises the ability of PDIA1 to refold HyPer_OX_, similarly to PDIA1 with no active site cysteines. For the refolding of roGFP2_OX_, the PDIA1*a*’_RED_ is even slightly more efficient than PDIA1 WT_RED_ and PDIA1*a*_RED_ is much less efficient (Figure 2C). It is clear from figures 2B-C that the major catalyst on reducing client substrates is the *a’* active site. Refolding catalysed by PDIA1*a*’_RED_ required reduction of the surrogate scrambled disulfide bond leading to the native state mostly reduced as indicated by the fluorescence intensity ratio (Figure 2D and 2E). If PDIA1_OX_ is added to the reaction mixture once a significant percentage of protein is refolded, dithiol oxidation is expected to occur based on the midpoint redox potentials of PDIA1 and native HyPer and roGFP2 discussed above and indeed, it is the case. Notably, PDIA1*a* is the major catalyst on the oxidation of HyPer_RED_ and roGFP2_RED_ after refolding confirming the greater susceptibility of the *a* domain towards DTT-mediated reduction *in situ* (Appenzeller-Herzog et al., 2010). Addition of the same PDIA1 mutants in the reduced state led to no protein oxidation as expected (see fade filled symbols in Figure 2D and 2E). Figure 2F summarizes the role of the *a’* active site of PDIA1 as the major catalyst for reduction and consequent refolding and the *a* active site as the main thiol oxidant, showing how scrambled disulfide bonds in proteins may be resolved through cycles of reduction-oxidation.

### Refolding and oxidation of HyPer and roGFP2 by PDIA1 and downstream oxidases

We tested the capability of PDIA1_RED_ to refold HyPer_OX_ in the presence of the glutathione couple GSH/GSSG. A total glutathione concentration of 15 mM and a ratio [GSH]/[GSSG] of 6.9 was used according to what was proposed to exist in the ER (Montero et al., 2013). PDIA1_RED_ is capable of refolding HyPer_OX_ in the presence of the glutathione couple with almost the same efficiency. Refolding was not significant (not kinetically competitive) in the presence of the glutathione couple only (left panel in figure 3A). Oxidised glutathione at mM concentration can promote the formation of the disulfide bond on HyPer once it was refolded (right panel in figure 3A).

**Figure 3.**
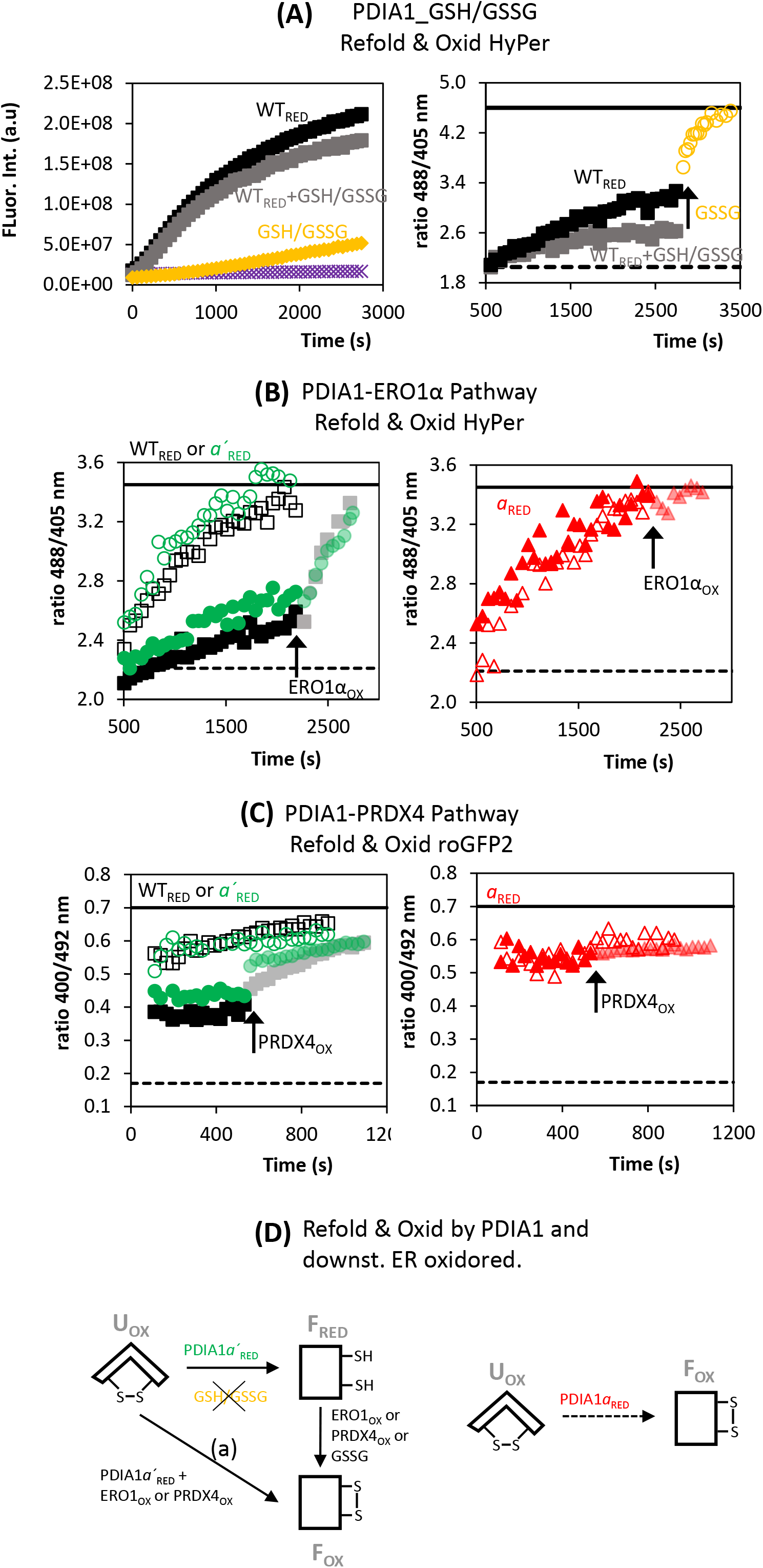
Panel (A) left panel shows the refolding of HyPer_OX_ measured from the time dependence of the integrated fluorescence intensity and right panel the degree of HyPer oxidation during refolding assessed through the ratio 488/405 nm. Reaction mixture contained PDIA1 WT_RED_ (■), PDIA1 WT_RED_ + 13.1 mM GSH + 1.9 mM GSSG (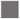), 13.1 mM GSH + 1.9 mM GSSG (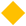) and buffer only (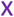). In the right panel, after refolding of HyPer_OX_ catalyzed by PDIA1 WT_RED_ only (■), 1.9 mM GSSG was added as depicted by the arrow (■ turn into 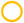). Dashed line at the bottom is the fluorescence ratio for the HyPer_RED_ standard and solid line at the top is the ratio for the HyPer_OX_ standard measured for this assay. Panel (B) shows the refolding of HyPer_OX_ and oxidation catalyzed by the reconstituted PDIA1-ERO1α pathway *in vitro*. Excitation fluorescence intensity ratio (488/405 nm) for the refolding of HyPer_OX_ catalyzed by PDIA1 WT_RED_ (■) and PDIA1*a*’_RED_ (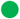) in the left panel and PDIA1*a*_RED_ (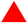) in the right panel. Refolding was recorded during the first 2200 s and then ERO1α_OX_ was added (pointed by arrow and filled symbols turned into fade filled symbols, same color). Empty symbols show the ratio 488/405 nm when refolding occurs in the presence of both PDIA1_RED_ and ERO1α_OX_ (catalyzed by PDIA1 WT_RED_ (□) or PDIA1*a*’_RED_ (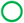) in the left panel and PDIA1*a*_RED_ (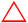) in the right panel). A control with ERO1α_OX_ only was carried out as well as with PDIA1 containing no active site cysteines but refolding was negligible and therefore no reliable values for the ratio 488/405 nm were measured (data not shown). Dashed line at the bottom is the ratio for the HyPer_RED_ standard and solid line at the top is the ratio for the HyPer_OX_ standard measured for this assay. Panel (C) Same legend as panel (B) but reporting the refolding of roGFP2OX during 600 s and oxidation (ratio 400/492 nm) catalyzed by the reconstituted PDIA1-PRDX4 pathway *in vitro*. Empty symbols show the ratio 400/492 nm when refolding occurs in the presence of both PDIA1_RED_ and PRDX4_OX_ and solid symbols for refolding in the presence of PDIA1_RED_ only. Dashed line at the bottom is the ratio for the roGFP2_RED_ standard and solid line at the top is the ratio for the roGFP2OX standard measured specifically for this assay. Panel (D): Schematic representation of the pathways for the refolding and oxidation of HyPer and roGFP2 catalyzed by PDIA1RED in the presence of the downstream oxidoreductases ERO1αOX and PRDX4_OX_. Left scheme for the PDIA1*a*’_RED_ and right scheme for PDIA1*a*’_RED_ (dashed arrow in right scheme means low refolding kinetics/efficiency). Pathway (a) in left scheme depicts the simultaneous addition of PDIA1*a*’_RED_ and oxidized downstream oxidoreductases where the native reduced state does not accumulate because kinetics of oxidation by ERO1α and PRDX4 are faster than refolding. U stands for the unfolded state and F for the folded and subscripts RED and OX for reduced and oxidised, respectively. Panels (A) to (C) show one representative assay from three independent experiments. PDIA1*a* (mutant C397S-C400S); PDIA1*a*’ (mutant C53S-C56S)

In the ER, besides PDIs other catalysts need to be involved in oxidative protein folding to keep the pool of PDI_OX_ equivalents and maintain sustained levels of disulfide bond formation in newly synthesized proteins. In the ER of mammals, two oxidoreductases, ERO1α and PRDX4 are responsible for recycling PDIA1 to become engaged in the oxidation of newly synthesised proteins (Zito et al., 2010; Tavender et al., 2010). We tested the refolding and oxidation of HyPer_OX_ catalysed by PDIA1_RED_ in the presence of ERO1α_OX_ (Figure 3B). If both, PDIA1 WT_RED_ and ERO1α_OX_ are added simultaneously to the reaction mixture, protein refolding is accompanied by dithiol oxidation as expected from the transfer of disulfide bonds from ERO1α to PDIA1 and to client protein (Figure 3B, empty black squares in left panel). This pattern was clear when refolding was catalysed by PDIA1 WT_RED_ or PDIA1*a’*RED both plotted in the left panel of figure 3B (empty black squares and empty green balls, respectively). In the absence of ERO1α during refolding much less oxidation was observed as expected, and posterior addition of ERO1α_OX_ catalysed disulfide bond formation (Figure 3B, filled symbols in left panel turned into fainted filled symbols, same colour, after addition of ERO1α_OX_ pointed by an arrow). Again, this pattern was clear for refolding catalysed by PDIA1 WT_RED_ and PDIA1*a*’_RED_. Differently, for PDIA1*a*_RED_ shown in the right panel of figure 3B, both refolding and oxidation occur even in the absence of ERO1α_OX_ and refolded HyPer ends up in the oxidised state with no further oxidation promoted by ERO1α_OX_ (Figure 3B right panel, compare empty and filled symbols). This distinct catalytic behaviour between the *a* and *a’* active site of PDIA1 is absolutely in accordance with: (i) the main role of the *a* active site is the oxidation of client proteins as shown previously in figure 2 and (ii) published data showed that electron transfer to ERO1α occurs mostly from the *a’* active site (Araki and Nagata, 2011; Chambers et al., 2010; Wang et al., 2009; Masui et al., 2011; Tsai and Rapoport, 2002). To summarize, refolding of HyPer_OX_ catalysed by PDIA1 WT_RED_ or by PDIA1*a*’RED ends up mostly with native reduced HyPer and disulfide bond formation on the client substrate required the presence of ERO1α_OX_ to pull electrons through the pathway. On contrary, refolding of HyPer_OX_ catalysed by PDIA1*a*_RED_ ends up with HyPer_OX_ independently of the presence of ERO1α. Electrons can flow from native client proteins to the *a* active site of PDIA1 for dithiol oxidation but cannot go further to ERO1α due to lack of electron transfer between the *a* active site and ERO1α. HyPer cannot be used as client protein to study the pathway PDI-PRDX4 as an external source of H_2_O_2_ is required to act as the final electron acceptor and HyPer is oxidised by nM levels of H_2_O_2_ (Melo et al., 2017). However, refolding and oxidation of roGFP2 can be studied in the presence of an exogenous source of H_2_O_2_ to fuel PRDX4 catalysis. When the downstream oxidoreductase is PRDX4 instead of ERO1α, both PDIA1 WT_RED_ and PDIA1*a*’_RED_ catalyse the refolding of roGFP2 and increased oxidation occurs if PRDX4_OX_ is present or added later (Figure 3C, compare empty symbols with filled symbols and fade filled symbols after later addition of PRDX4_OX_ in the left panel). For PDIA1*a*_RED_, roGFP2 refolded was mostly oxidised and the presence or later addition of PRDX4_OX_ does not promote additional dithiol oxidation (Figure 3C, right panel). As shown above for ERO1α, the *a*’ active site of PDIA1 is involved in transferring electrons to PRDX4, confirming previous observations (Tavender at al., 2010). RoGFP2 oxidation increases with the simultaneous or later addition of PRDX4_OX_ only if PDIA1 WT or PDIA1*a*’ were the catalysts. No further dithiol oxidation was observed with PRDX4_OX_ addition if PDIA1*a* was used as catalyst instead. Figure 3D summarizes our observations for the refolding and oxidation of HyPer and roGFP2 when the enzymatic pathways that catalyse oxidative protein folding in the ER were reconstituted *in vitro*. The PDIA1*a*’ is more efficient in thiol reduction and refolding, and catalysis of disulfide bond in the refolded protein requires the presence of one of the downstream oxidoreductases ERO1α or PRDx4. PDIA1*a* is much less efficient in the refolding of client proteins (because is more oxidising) but the lower levels of refolded protein are promptly oxidised with no electrons flowing to downstream oxidoreductases.

### Oxidation of native HyPer and roGFP2 by PDIA1 and downstream oxidases

Besides refolding of HyPer and roGFP2, we also tested oxidation of both proteins in the native state by the two enzymatic pathways that operate in the ER. Native HyPer_RED_ and roGFP2_RED_ were the substrates and PDIA1 in the reduced state (to prevent substrate oxidation by PDIA1 alone) plus ERO1α_OX_ were the catalysts. In the presence of ERO1α_OX_, both PDIA1 WT and PDIA1*a’* were able to oxidize HyPer and roGFP2 as client proteins with close efficiencies (Figure 4A and 4B). On contrary, PDIA1*a* along with ERO1α_OX_ was unable to significantly oxidize client substrates, showing once more that ERO1α_OX_ can transfer disulfide bonds to the *a’* active site of PDIA1 only. The same behavior was observed when the PDIA1-PRDX4 pathway is reconstituted to oxidize roGFP2_RED_ (Figure 4C). Oxidation was observed in the presence of PDIA1 WT and PDIA1*a’* but not for PDIA1*a*. Preferential transfer of disulfide bonds from ERO1α and PRDX4 to the *a*’ active site of PDIA1 was previously reported (Araki and Nagata, 2011; Chambers et al., 2010; Wang et al., 2009; Masui et al., 2011; Tavender et al., 2010; Tsai and Rapoport, 2002). Oxidation of native HyPer and native roGFP2 by the two pathways PDIA1-ERO1α and PDIA1-PRDX4 is channeled almost exclusively through the *a*’ active site of PDIA1 (Figure 4D).

**Figure 4.**
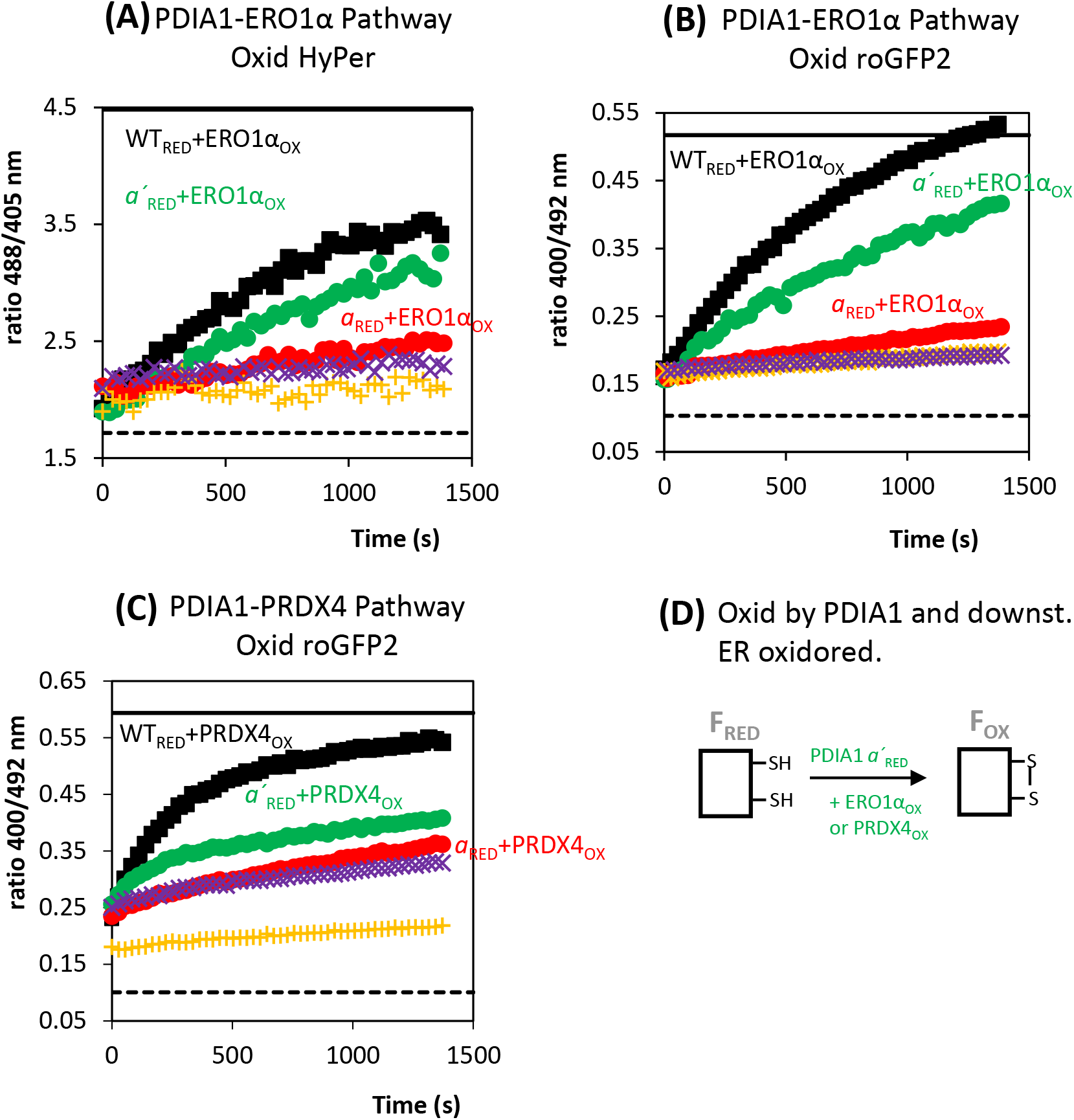
Panel (A) Oxidation of native HyPer_RED_ by the reconstituted PDIA1-ERO1α pathway *in vitro*. Excitation fluorescence intensity ratio (488/405 nm) for the oxidation of HyPer_RED_ catalyzed by PDIA1 WT_RED_ plus ERO1α_OX_ (■), PDIA1*a*_RED_ plus ERO1α_OX_ (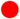), PDIA1*a*’_RED_ plus ERO1α_OX_ (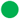), PDIA1 WT_RED_ in the absence of ERO1α (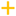) and ERO1α_OX_ only (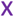). Panel (B) Same legend as panel (A) but reporting the oxidation (ratio 400/492 nm) of roGFP2_RED_ by the reconstituted PDIA1-ERO1α pathway *in vitro*. Panel (C) Same legend as panel (A) but reporting the oxidation (ratio 400/492 nm) of roGFP2_RED_ by the reconstituted PDIA1-PRDX4 pathway *in vitro*. The presence of glucose oxidase and glucose as an exogenous source of H_2_O_2_ to act as the final electron acceptor for PRDX4 were required for this assay. Symbols (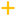) refer to the absence of PRDX4_OX_ and presence of PDIA1 WT_RED_ and symbols (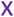) to the presence of PRDX4_OX_ plus glucose and glucose oxidase. In all panels, the dashed line at the bottom is the ratio for HyPer_RED_ or roGFP2_RED_ standards and solid line at the top is the ratio for HyPer_OX_ or roGFP2_OX_ standards. Panel (D): Schematic representation for the oxidation of HyPer and roGFP2 catalyzed by PDIA1*a*’_RED_ and downstream ER oxidoreductases ERO1_OX_ or PRDX4_OX_. PDIA1 was in the reduced state to prevent HyPer/roGFP2 oxidation by PDIA1 before its oxidation by downstream oxidoreductases. F stands for the folded state and subscripts RED and OX for reduced and oxidized, respectively. Panels (A) to (C) show one representative assay from three independent experiments. PDIA1*a* (mutant C397S-C400S); PDIA1*a*’ (mutant C53S-C56S)

### Disulfide bond between the two active sites and compactness of PDIA1

It has been shown that PDIA1 can adopt at least two conformers with a different distance between the two arms containing the catalytic sites (Nguyen et al., 2008; Inagaki et al., 2015). X-ray crystallography has shown that in oxidised PDIA1 the two active sites are 40.3 Å apart (Protein Data Bank 4EL1) and in the reduced form (PDB 4EKZ) the *a’* domain twists around 45°, approaching the two active sites to 27.6 Å (Wang et al., 2013). Moreover, *in silico* studies showed that the two catalytic domains can locate much closer, up to a distance compatible with a disulfide bond between the two catalytic sites (Yang et al., 2014). Mutagenizing the two resolving cysteines into serines (C56S-C400S) leads to the formation of a stable disulfide bond between attacking cysteines of the two active sites as shown in a non-reducing gel (Figure 5A and Yang et al., 2014). The two arms of the PDIA1 horseshoe shape are flexible enough to approach the two active sites to a distance that allows the formation of a disulfide bond between them. It should be transient in the WT but becomes stable in the absence of resolving cysteines. An interdomain disulfide bond between the two active sites should lead to a more compact PDIA1 conformation and to address this issue we have used the switch sense biophysical analysis (Langer et al., 2013). A DNA double strand nanolever with 48bp immobilized on a chip, moves upward, away from the surface of the chip, when a negative electric potential is applied to the chip due to the negative charge of DNA. If the tip of the nanolever has a fluorophore attached, its fluorescence will increase during the upward motion away from the quenching effect of the gold surface of the chip. The upward movement of the nanolever can be followed over 10 μs time through the increase in fluorescence intensity (Figure 5B). The complementary DNA strand on the nanolever can be derivatized with a NTA group to immobilize PDIA1 through the His-tag. Binding PDIA1 to the nanolever slows down the upward movement compared to the NTA group only. Slowed molecular dynamics of the nanolever depend on the size and compactness of PDIA1 (Langer et al., 2014). Binding of PDIA1 with no resolving cysteines, results in a faster motion upward compared with PDIA1 WT, reflecting the greater compactness of the protein with a disulfide bond between the two active sites. Interestingly, PDIA1 with no cysteines in the active sites is as compact as PDIA1 with the disulfide bond connecting the two active sites showing that the average distance between the active sites located in opposite arms of PDIA1 depends on the presence of cysteines in the active sites (Figure 5B). Dynamics of the upward movement were equated to the area under the fluorescence curve, the so-called dynamic response, which is larger for more compact protein conformers as it is the case of PDIA1 with a disulfide bond between the two active sites and with no cysteines in the active sites (Figure 5C).

**Figure 5.**
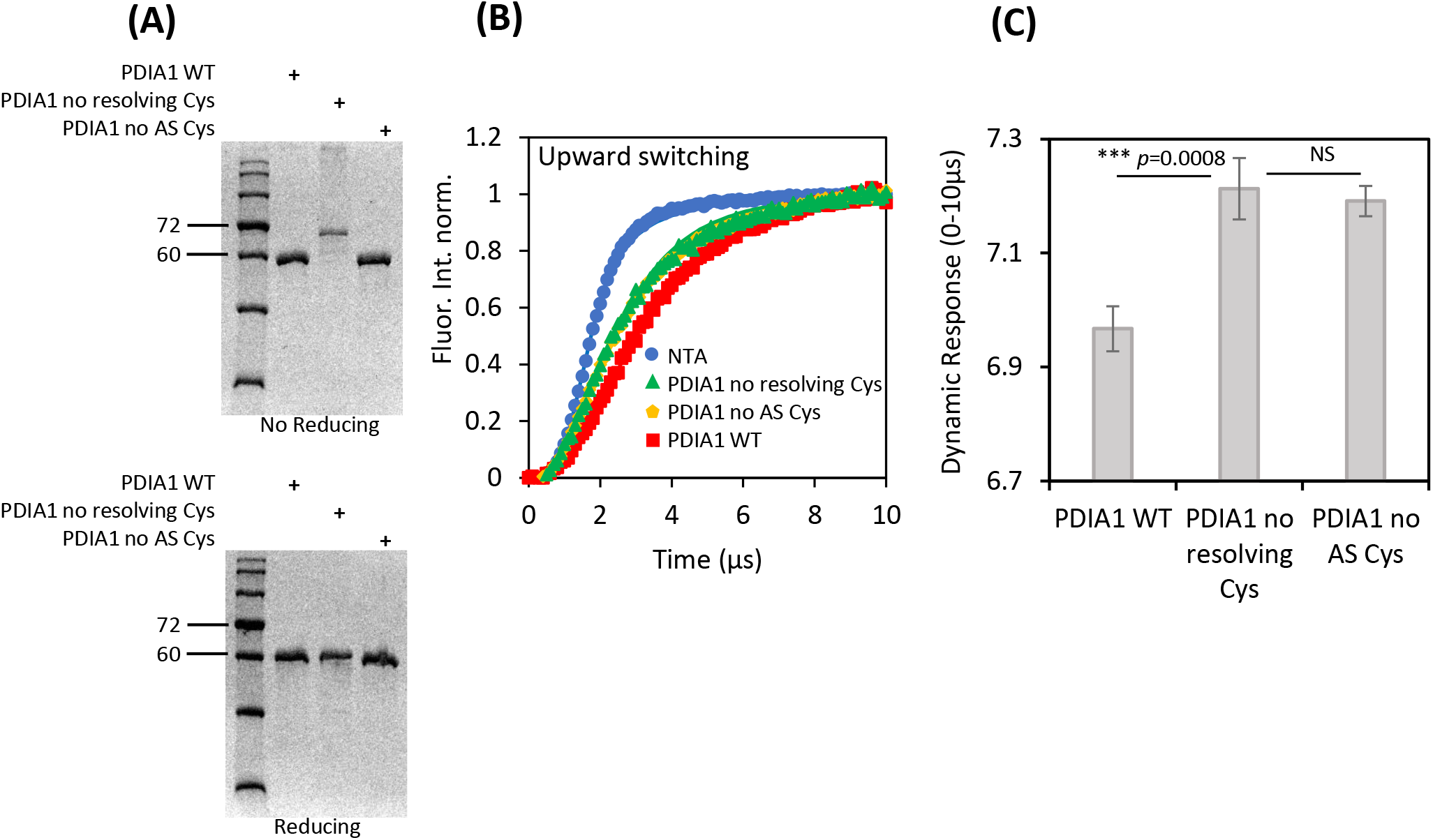
Panel (A) No reducing and reducing SDS-Page gels of PDIA1 WT, PDIA1 with no resolving cysteines (PDIA1 C56S-C400S) and with no active site (AS) cysteines (PDIA1 C53S-C56S-C397S-C400S). PDIA1 MW is ~57 kDa. Panel (B) Electro-switchable DNA double-stranded nanolevers derivatized at the DNA’s top end with nitriloacetic acid (NTA) during its movement upward from the surface of the biochip after applying a repulsive potential of 10 kHz. An epifluorescence setup measured the increase in fluorescence from the fluorophore attached to the NTA-complementary DNA strand during nanolever upward motion (more technical details in material and methods and in Langer at al., 2013). DNA nanolever with NTA only (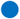) moves faster than after binding PDIA1 WT (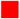) through the Histag which increased hydrodynamic friction during upward motion (longer times are needed to reach maximum fluorescence when the DNA nanolever tilt away from the surface). More compact conformations than PDIA1 WT such as PDIA1 with no resolving cysteines (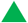) or with no cysteines in the active site (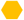) cause less friction reaching maximum fluorescence in shorter times. Panel (C) Dynamic response equating to the area under the fluorescence curve when switching upward (corresponding to the change in height in fluorescence units that is covered by the top end of the DNA nanolever within a given time period, 0-10 μs) of PDIA1 WT, PDIA1 with no resolving cysteines and PDIA1 with no active site cysteines for 18 measurements as in panel (B). Larger dynamic responses result from more compact PDIA1 conformations. One-way ANOVA test for independent measurements, \*\*\**p*<0.001, NS not significant.

### Catalytic cooperativity between the two active sites of PDIA1 on the oxidation of native roGFP2 by the PDIA1-ERO1α pathway

Catalytic cooperativity between the two active sites of PDIA1 on thiol oxidation was reported (Araki and Nagata, 2011; Chambers et al., 2010). Using native roGFP2_RED_ as substrate, we have compared the oxidative catalytic rates of increasing concentrations of PDIA1_RED_ in the presence of ERO1α_OX_ (Figure 6A). Substrate oxidation may occur only if the full pathway PDIA1-ERO1α is active as PDIA1 was added to the reaction in the reduced state. As expected, no oxidation was observed for PDIA1*a* as it cannot transfer electrons to ERO1α. Cooperativity between the two active sites leads to larger catalytic rates of PDIA1 WT compared to PDIA1*a*’, reflecting the contribution of the *a* active site to roGFP2 oxidation in the context of the full pathway. As proposed elsewhere, this increased catalytic rate for the WT can be rationalised if electrons flow from *a* to *a’*, with the latter being the preferential path to transfer electrons to ERO1α (Araki and Nagata, 2011). In the crystal, the distance between the two active sites (changes from 40.3 to 27.6 Å upon reduction) is not compatible with the formation of a transient interdomain disulfide bond between them (average length for a reversible disulfide bond is 2.18 Å (Sun et al., 2017)) but relying on PDIA1 conformational flexibility that can occur in solution (Figure 5A). The redox-regulated conformational change that brings the *a’* active site closer to *a* is impaired if the cation-π interaction between R300 and W396 is broken (Wang et al., 2013). Therefore, we have investigated if the cooperativity between the two active sites of PDIA1 is abolished for the PDIA1 mutant R300A, and indeed it was (Figure 6A). In the R300A mutant, the *a* active site no longer contributes to the oxidation of roGFP2 in the context of the full pathway despite being more oxidising than *a’*. Electrons no longer flow from the *a* to ERO1α through the *a’* active site when the main interaction that brings the two active sites closer was disrupted. Decreased catalytic rates observed for the R300A mutant in comparison to the WT cannot be assigned to structural changes resulting from the mutation as both the tertiary and secondary structure of R300A were not different from those of the WT (Figure 6B and 6C). Cooperativity between the two active sites of PDIA1 is also abolished for the W396A PDIA1 mutant (Figure S6), although in this case might be argue that the vicinity of W396 to the catalytic cysteines of the *a’* active site (C397 and C400) may affect its redox potential.

**Figure 6.**
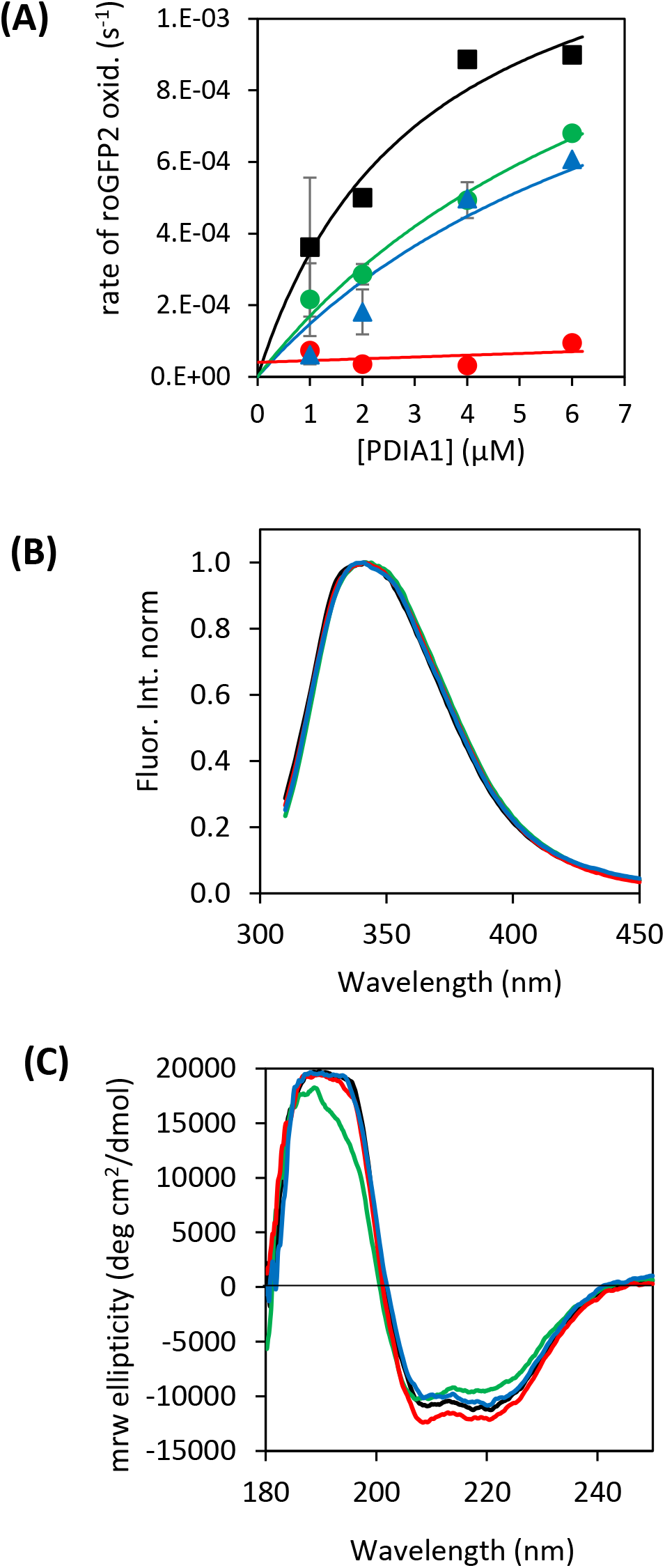
Panel (A) Rate of roGFP2 oxidation by increasing concentrations of PDIA1 WT and mutants (WT_RED_ ■; *a*’_RED_ 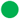; *a*_RED_ 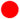; R300A_RED_ 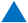) plus 3 μM ERO1αOX. PDIA1 was added reduced to prevent roGFP2 oxidation unless PDIA1 is oxidised by ERO1α. Panel (B) and (C) show tryptophan fluorescence emission and far-UV CD spectra of PDIA1 WT and mutants to probe tertiary and secondary structure, respectively (WT, black; *a*’, green; *a*, red; R300A blue).

## Discussion

RoGFP2 and HyPer are fluorescent proteins engineered with cysteine residues to sense redox poise. The introduction of these cysteines and their engagement in a disulfide bond make these proteins suitable substrates to measure the activity of thiol-disulfide oxidoreductases such as PDIA1. Moreover, they also revealed to be substrates for the isomerase activity of PDIA1. Once they are unfolded in the oxidised state by chemical they cannot refold back to the native state. The engineered disulfide bond acts like a cufflink that traps the protein conformation in a compact misfolded state with no fluorescence in the visible range (Figure 1 and Correia et al., 2020). To resolve this trap and allow the protein to refold, the engineered disulfide bond needs to be reduced. This can be achieved with a reducing agent at mM concentrations or even more efficiently by catalytic μM amounts of PDIA1 reduced (Figure 2). PDIA1 in the reduced state but not in the oxidised state, can reduce the disulfide bond and assist in protein refolding through its chaperone activity (Cai et al., 1994; Song and Wang, 1995). Based on the redox potentials (−260 mV for HyPer, −272/−280 mV for roGFP2 and −206 mV for PDIA1), PDIA1 was expected to catalyse disulfide bond formation on HyPer and roGFP2. However, reduction of client proteins by PDIA1 during refolding becomes possible if the redox potential of the sensing cysteines in the misfolded protein is more oxidising than in the native protein. Non-native disulphide bonds are known to form in proteins under normal physiological conditions (Jansens et al., 2002). These scrambled disulfide bonds may be resolved in cycles of reduction-oxidation or by direct isomerization (Hatahet and Ruddock, 2019). In a cycle of reduction-oxidation, PDIA1 reduced can reduce the disulfide bond with higher redox potential in the compact misfolded client and then PDIA1 oxidised can form the native disulfide bond on the client protein in accordance with the redox potentials stated above (Figure 2).

While PDIA1*a’* is significantly more efficient on the reduction of the disulfide bond in the compact misfolded substrate to promote refolding, PDIA1*a* is more efficient on catalysing the disulfide bond once HyPer and roGFP2 are refolded (Figure 2). Moreover, refolding induced by PDIA1*a’* results in a native protein mostly reduced while refolding induced by PDIA1*a*, despite significantly less efficient, results in larger degree of substrate oxidation (Figure 3). In our refolding/oxidation assays with two different client proteins, there is a clear distinction for the role assigned to the two active sites of PDIA1 despite the same redox potential reported for both (Bekendam et al., 2016). The different role assigned for the two active sites, *a* more oxidising and *a’* more reducing, can be rationalised if the redox potential of the active site cysteines shifts with PDIA1 conformation and interaction with clients (Hatahet and Ruddock, 2009). Indeed, binding of small ligands induces movement of the x-linker leading to a more compact PDIA1 conformation and shifts the active-sites to a more reduced state (Bekendam et al., 2016).

Two oxidoreductases present in the ER of mammals, ERO1α and PRDX4, recycle PDIA1 to become engaged in the oxidation of newly synthesised proteins (Zito et al., 2010; Tavender et al., 2010). It was shown that the *a’* active site of PDIA1 is the one that preferentially transfer electrons to ERO1α (Araki and Nagata, 2011; Chambers et al., 2010; Wang et al., 2009; Masui et al., 2011; Tsai and Rapoport, 2002) and to PRDX4 (Tavender et al., 2010). Refolding and oxidation of HyPer and roGFP2 *in vitro* by the full pathways operating in the ER, confirmed the almost exclusive involvement of the *a’* active site on the transfer of electrons to ERO1α and PRDX4 (Figure 3). To uncouple any effect that may result from protein refolding we also tested native HyPer and roGFP2 as client substrates for the full pathways operating in the ER. Again, ERO1α and PRDX4 oxidise almost exclusively the *a’* active site of PDIA1 (Figure 4).

While the catalytic activity of domains within PDIA1 are capable of autonomous activity, they demonstrate cooperative and enhanced behaviour within the context of the WT (Araki and Nagata, 2011; Chambers et al., 2010; Hatahet and Ruddock, 2009). The mechanism of this cooperativity, however, is poorly understood particularly in the context of the full pathways operating in the ER. In the context of the full pathways PDI-ERO1α and PDI-PRDX4, no oxidation of HyPer and roGFP2 was observed when the catalyst is PDIA1*a* (Figure 4 and 6). Nevertheless, cooperativity between the two active sites results in a faster oxidative rate of the WT compared to PDIA1*a’* as we have shown and reported before in the presence of ERO1α (Araki and Nagata, 2011; Chambers et al., 2010). How can the *a* active site contribute to the activity of PDIA1 WT if its oxidative capacity in our assays depends on the presence of a downstream oxidase that receives electrons from the *a’* active site? Does it seem rational that PDIA1 has evolved to have the more oxidising *a* active site (Figure 2) unfitted to transfer electrons to any of the two downstream oxidoreductases involved in oxidative protein folding in the ER (Figure 3 and 4)? Experimental evidence for disulfide exchange between the two active sites was gathered previously (Chambers et al., 2010; Araki and Nagata, 2011; Freedman et al., 2017; Yang et al., 2014; Araki et al., 2013) but the functional implications in the context of the different PDIA1 activities were not clear. Our results indicate an interdomain disulfide exchange as the mechanism to solve the paradoxes stated above. We showed that the conformational flexibility of PDIA1 allows the formation of a disulfide bond between the catalytic sites *a* and *a’* (Figure 5A). This interdomain disulfide bond is stable only if the two resolving cysteines (C56 and C400) are absent, but its formation shows that the conformational flexibility allows the approximation between the two arms of the horseshoe shape of PDIA1 to a distance compatible with the formation of a disulfide bond. Cross-linking assays also showed that in solution, the two active sites can approach much nearer than the distances measured in the crystal (Hawkins et al., 1991; Peng et al., 2014) and *in silico* flexibility modelling and molecular dynamics showed hinge-bending of domains *a* and *a’* to close contact (Freedman et al., 2017), enough for an interdomain disulfide bond (Yang et al., 2014). Formation of this interdomain disulfide bond, results in a more compact PDIA1 conformation compared to the WT (Figure 5B and 5C). Interestingly, in the absence of any active site cysteine, PDIA1 is as compact as the interdomain disulfide bonded state, highlighting the role of active site cysteines on the dynamics that change the distance between the two active sites. Indeed, this distance was shown to be redox-dependent with an interdomain conformational change moving the *a’* domain closer to *a* when the active site cysteines are reduced (Wang et al., 2013; Inagaki et al., 2015). Figure 7 schematises the role of disulfide exchange between the two active sites as an interdomain redox relay that explains the role of each active site, the cooperativity between them and the interplay with downstream oxidases. The *a* active site is the major player on the oxidation of cysteine residues on client proteins. Reducing equivalents are then transferred from the *a* to the *a’* active site (interdomain redox relay), explaining the cooperativity between the two active sites on oxidative protein folding. This interdomain redox relay requires conformational plasticity and a change in the redox potential of catalytic cysteines which should result from the conformational change that leads to a more compact PDIA1 conformation. The reduced *a’* active site can either be involved in the reduction of client proteins (resolving the misfolded compact state of HyPer and roGFP2 as shown in figure 2) or transfer the reducing equivalents to the downstream oxidases ERO1α or PRDX4, a process occurring mostly from the *a’* active site. Recycling of the oxidised state of the *a’* active site is thus possible through reduction of scrambled disulfide bonds or through oxidation by downstream oxidases, depending on the encounter rate, affinity, and redox potential of scrambled disulfide bonds which should be more oxidising than native disulfide bonds as documented for HyPer and roGFP2. This path for electron transfer resolves scrambled disulfide bonds through reduction-oxidation cycles, which seems to be the predominant reaction mechanism (Schwaller et al., 2003), but does not compromise the possibility of direct isomerization of scrambled disulfides with no net change in the redox state of the *a’* active site (Hatahet and Ruddock, 2009). Consistent with the interdomain redox relay mechanism, the redox state of PDIA1 in cells can be fully oxidised, semi-oxidised (either *a* or *a’* oxidised) or fully reduced when reducing equivalents overwhelmed PDIA1 electron transfer capacity (Appenzeller-Herzog and Ellgaard, 2008). Yeast PDI crystallographic structure showed the two active site cysteines in the *a* domain primarily in the oxidised state, while their counterparts in the *a’* domain are in the reduced state (Tian et al., 2006). In addition, the open-closed conformational change of PDIA1 underlying the interdomain redox relay, depends on the redox-state of the a’ domain (Inagaki et al., 2015; Wang et al., 2013). The interdomain redox relay mechanism proposed here also rationalises why ERO1α activity is switched off when reduced PDIA1 is depleted (Appenzeller-Herzog et al., 2008): To pile up more reducing equivalents in the *a’* active site needed for the reduction of non-native disulfide bonds. Redox regulation of ERO1α activity by the a’ domain of PDIA1 (Kanemura et al., 2016) should be linked to the necessity of keeping the pool of reducing equivalents on PDIA1 rather than preventing excessive formation of H_2_O_2_ which may be reduced by PRDX4 (Zito et al., 2010; Tavender et al., 2010), glutathione peroxidases (Kanemura 2020a; Nguyen et al., 2011) or buffered by glutathione above a critical threshold concentration (Melo et al., 2017). Another observation in line with the mechanism of figure 7 is the inactivation of the pathway PDIA1-ERO1α when levels of unfolded proteins and folding intermediates exceed the folding capacity of the system (Moilanen and Ruddock, 2020). Binding of PDIA1 to non-native proteins seems to compete with binding to ERO1α inhibiting its activation to keep unfolded proteins and folding intermediates mainly in the reduced state. This mechanism may be the basis of the effective proof reading of non-native disulfide bonds by PDIA1 compared to others PDIs such as ERp46 and P5 (Sato et al., 2013). We propose the interdomain redox relay mechanism to be the preferential pathway for electron transfer through PDIA1, although some promiscuity between the two active sites of PDIA1 may exist. We have observed slow kinetics of PDIA1*a* on reducing the scrambled disulfide bond on client proteins, slow kinetics of PDIA1*a’* on oxidising native thiols and very slow kinetics for the transfer of electrons from PDIA1*a* to downstream oxidases (Figure 2 and 4). However, this promiscuity might be irrelevant in the ER where kinetics should determine which pathways go forward.

**Figure 7.**
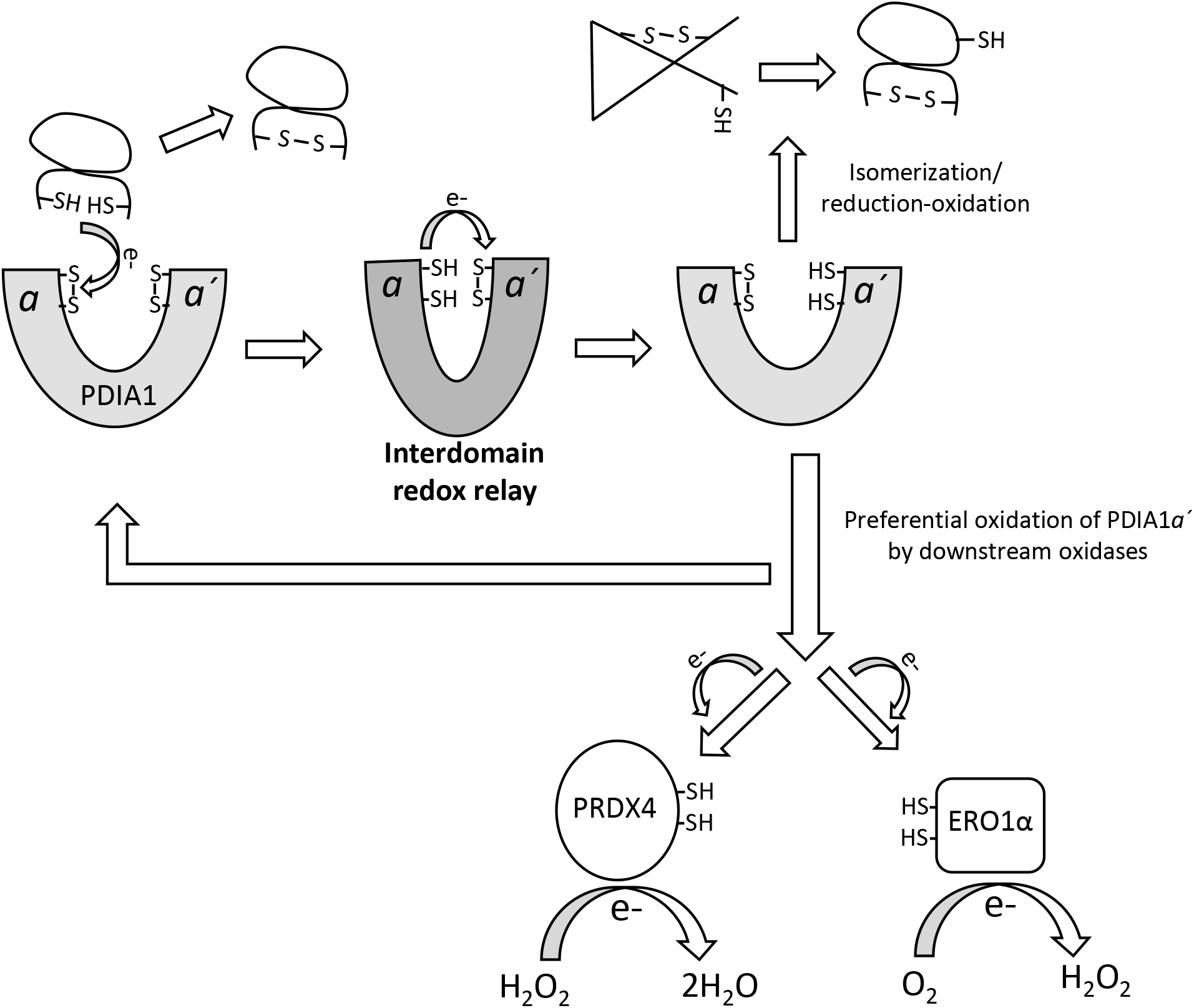
Mechanistic representation for the involvement of the interdomain redox relay of PDIA1 in oxidative protein folding. The PDIA1*a* is the major player on the oxidation of cysteine residues on client proteins (Fig. 2D, E). Reducing equivalents are then transferred from the *a* to the *a’ active site* via the interdomain redox relay in the closed conformation of PDIA1. The reduced *a*’ active site (more reducing, Fig. 2B, C) can either be involved in the reduction/isomerization of client proteins or in the transfer of reducing equivalents to downstream oxidases ERO1α or PRDX4 as the *a’* active site is the preferential path to communicate with downstream oxidases (Fig. 3B, C; Fig. 4A, B, C). If isomerization proceeds via cycles of reduction-oxidation (Karala et al., 2010), reduction of scrambled disulfides on client proteins is preferentially catalysed by the *a’* active site. Recycling of the oxidised state of the *a’ active site* can then be achieved through reduction of scrambled disulfide bonds or through oxidation by downstream oxidases depending on the encounter rate, affinity, and redox potential of scrambled disulfide bonds which are more oxidising than native disulfide bonds as observed for misfolded roGFP2 and HyPer.

A direct role of glutathione on the formation of correct disulfide bonds was challenged (Tsunoda et al., 2014) and changes in the concentration of glutathione in cells does not seem to affect the redox state of PDIA1 (Chakravarthi and Bulleid, 2004). We have observed that the pair GSH/GSSG, at concentrations reported to be present in the ER (Montero et al., 2013), had no significant effect on the refolding and oxidation of HyPer during the time scale of the assay, indicating that reduction/oxidation of client proteins by glutathione is not kinetically competitive with enzymatic machinery (Figure 3A). For the interdomain redox relay mechanism proposed, glutathione is not required for the modulation of the redox state of PDIA1. The pathway is sustainable if input (from client proteins) and output of reducing equivalents (for downstream oxidases and non-native disulfide bonds) match. In accordance, GSH depletion had no consequences for the formation of correct disulfides in proteins whose maturation or degradation require the reduction of non-native disulfide bonds (Tsunoda et al., 2014). A role of glutathione in the redox state of other ER oxidoreductases cannot be ruled out as shown for the reduction of PDIA3, also known as ERp57 (Jessop and Bulleid, 2004). Anyway, the main role of the glutathione couple in the ER seems to be providing buffering capacity against both oxidative and reductive stress. It plays an important role on the elimination of H_2_O_2_ from the ER under oxidative stress, i.e., when H_2_O_2_ levels increase above a critical threshold (Melo et al., 2017).

One alternative mechanism to the interdomain redox relay proposed is the intermolecular electron transfer from the *a* domain of one PDIA1 to the *a’* domain of other PDIA1 molecule. Oxidised PDIA1 was shown to form a face-to-face homodimer induced by unfolded substrates that creates a central hydrophobic cavity with multiple redox-active sites (Okumura et al., 2019). However, intermolecular electron transfer will increase the cooperativity in the WT at increased concentrations of PDIA1, which was not observed in our study (Figure 6A). Also, a mixture of PDIA1 molecules with the *a* or the *a’* active site does not display cooperativity, contradicting the intermolecular electron transfer mechanism (Araki and Nagata, 2011). Oxidised PDIA1 dimers are weak and transient and were proposed to be involved in the introduction of disulfide bonds into highly flexible unfolded substrates (Okumura et al., 2019).

The interdomain redox relay mechanism relies on the dynamics of PDIA1 conformation approaching the two active sites. An interaction between R300 and W396 was identified to stabilizing the compact conformation of PDIA1 with the active sites closer (Wang et al., 2013) and the mutant R300A can no longer maintain a closed conformation (Okumura et al., 2019). Removal of this interaction (for the R300A and W396A PDIA1 single mutants) abolished the cooperativity between the two active sites, an observation in strong support of the interdomain redox relay mechanism (Figure 6A and S6). No structural changes that may compromise PDIA1 activity were detected for the R300A and W396A mutants (Figure 6B and 6C), proving that cooperativity is totally dependent of the stabilization of the conformation with the two active sites in closer proximity. The physiological relevance of electron transfer between the two active sites at the cellular and organism level, was revealed by the mutation R300H in PDIA1 which was linked to amyotrophic lateral sclerosis (Woehlbier et al., 2016). Extensive evidence for the conformational flexibility and movement between the catalytic domains *a* and *a’* supporting the interdomain redox relay mechanism has been gathered over the years (Freedman et al., 2017). The orientation of PDIA1 domains change upon binding to large protein substrates with the movement of the *a’* domain being the most pronounced change (Biterova et al., 2019). A closed and open conformation of PDIA1 seem to be part of the catalytic cycle representing a substrate -bound and substrate-free states (Wang et al., 2013). Moreover, in silico analysis suggests that interconversion between the closed and open states is possible through domain motion with great variation of the distance between the two active sites (Freedman et al., 2017). An allosteric switch caused by binding of small molecules to the substrate-binding pocket on the *b’* domain modulates the catalytic activity of PDIA1 by increasing the reductase activity of the *a’* domain (Bekendam et al., 2016). More recently, single-molecule analysis revealed that oxidized PDIA1 is in rapid-equilibrium between open and closed conformations (Okumura et al., 2019).

In conclusion, the interdomain redox relay at the core of PDIA1 activity solves the conundrum of having the *a* active site more oxidising but unfitted to transfer electrons to downstream ER oxidases, explains the capability of PDIA1 to act as thiol oxidant forming native disulfides or as reductant to resolve scrambled disulfides and proposes a new rational for shutting down oxidative protein folding in the ER under redox imbalance or when the levels of unfolded proteins and folding intermediates exceed the folding capacity of the system.

## Acknowledgements

This study received Portuguese national funds from FCT - Foundation for Science and Technology through project UIDB/04326/2020, UIDP/04326/2020 and LA/P/0101/2020, and from the operational programmes CRESC Algarve 2020 and COMPETE 2020 through project EMBRC.PT ALG-01-0145-FEDER-022121.

