## Supplementary Figures and Table for "A conformational-dependent interdomain redox relay at the core of Protein Disulfide Isomerase activity"

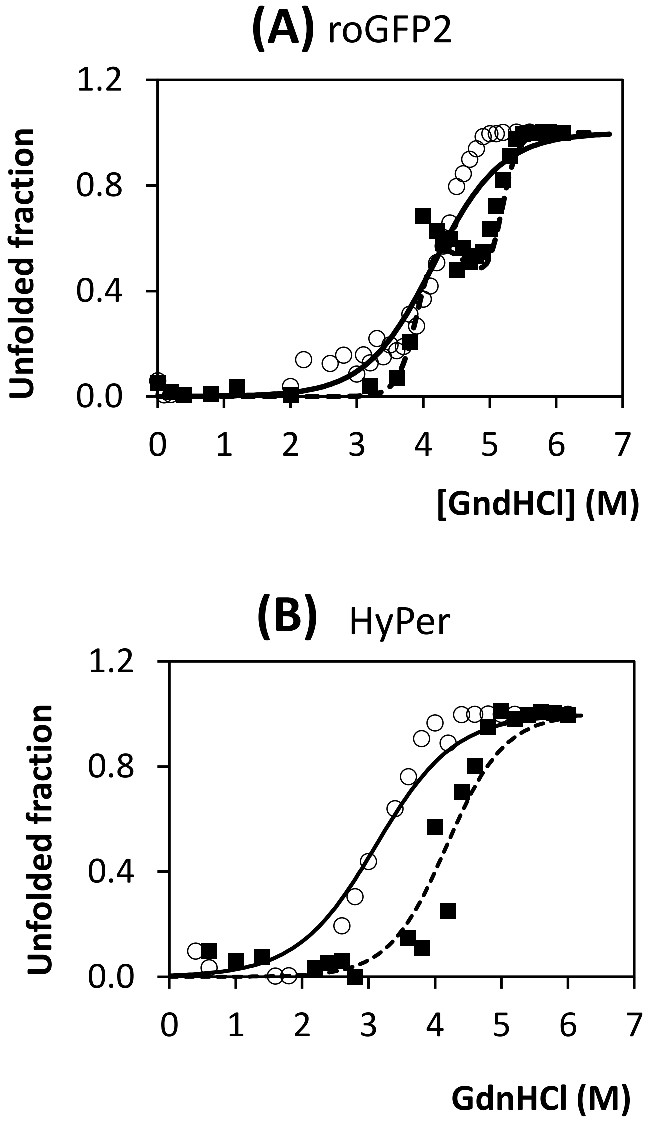


Figure S1. Panel (A): Unfolding of roGFP2 reduced in the presence of 20mM DTT (○) and oxidized (■) induced by GdnHCl and quantified through the integration of the excitation spectra. The solid line is the fit of the unfolding profile according to a two-state process (only the folded and unfolded states are present) and the dashed line is the fit for the unfolding of oxidized roGFP2 where two transitions had to be considered. The presence of the disulfide bond at the surface of the β-barrel structure of roGFP2 changes the unfolding pathway leading to the accumulation of an intermediate state. The stability increases from 8.1 to 34.5 kcal/mol and the mid-point of unfolding increases from 4.2 to 4.6 M of GdnHCl upon oxidation showing that the disulfide bond stabilizes the folded state relatively to the unfolded state. Panel (B): Unfolding of HyPer reduced in the presence of 20mM DTT (○) and oxidised (■) induced by GdnHCl. The following stability parameters were obtained for HyPer: $\Delta G^{water}$ 5.2 and 9.5 kcal/mol and mid-point of unfolding 3.1 and 4.2 M GdnHCl for the reduced and oxidised species, respectively. Therefore, contribution of the disulfide bond to the stability of HyPer is 4.3 kcal/mol. The solid and dashed lines are the fits of the unfolding profile according to a two-state process.

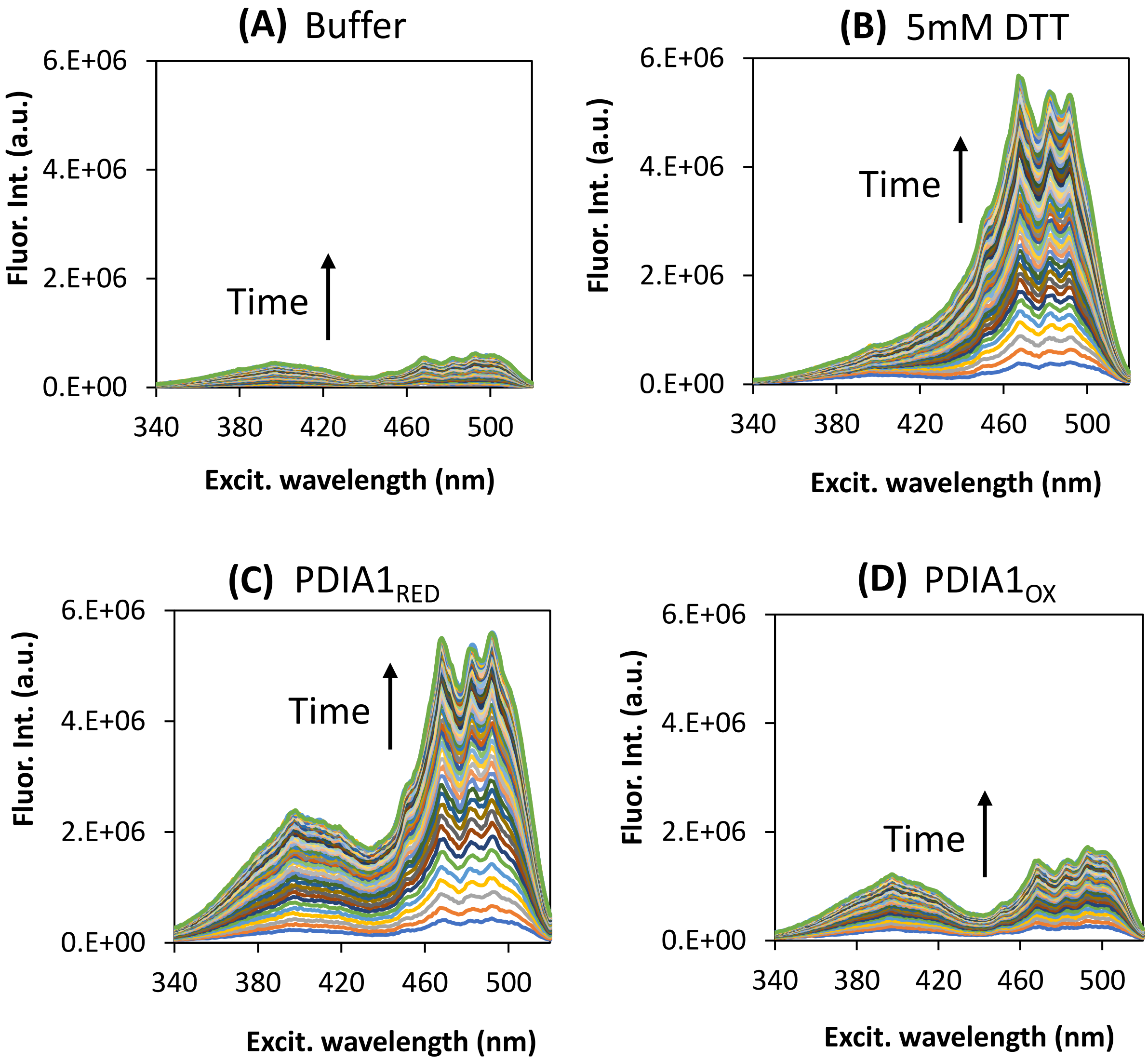


Figure S2. Refolding of oxidised chemically unfolded roGFP2. Unfolding was achieved at 5.4 M GdnHCl and refolding promoted by dilution to 0.2 M. Refolding and concomitant fluorescence emission does not occur in buffer (panel A) but DTT at 5 mM can reduce the disulfide bond that prevents refolding and the native reduced state with a fluorescence intensity ratio 400/492 nm of 0.21 is reached (panel B). Significant refolding was observed in the presence of 5 µM PDIA1 reduced (panel C) with the final state being partly oxidised with a ratio 400/492 nm of 0.41 compared to 0.71 for fully oxidised roGFP2. No significant refolding occurred in the presence of 5 µM PDIA1 oxidised (panel D).

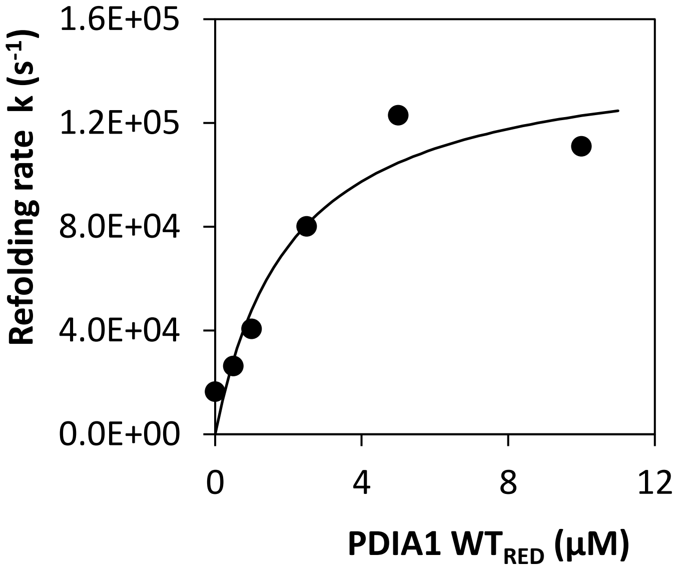


Figure S3. Rate of refolding of HyPer oxidised catalyzed by different concentrations of reduced PDIA1 WT obeys a hyperbolic equation as expected from enzyme kinetics (data calculated from the initial velocities of HyPer refolding profiles shown in figure 2A)

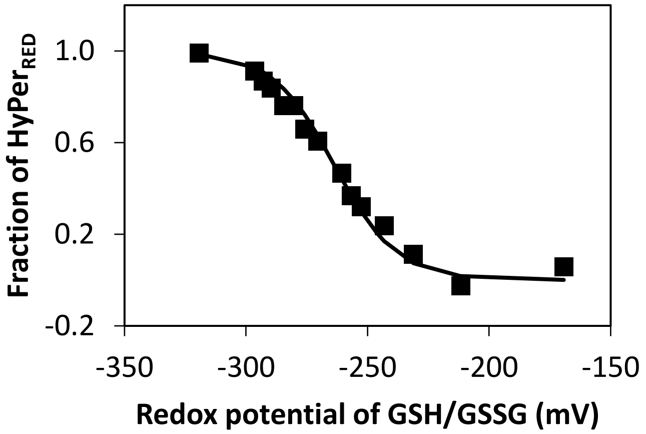


Figure S4. Redox equilibrium of HyPer upon titration with the redox couple GSH/GSSG. The midpoint potential ($E_0´$) of HyPer was determined as -260±5 mV by titration against GSH/GSSG mixtures using the Nernst equation according to Hanson et al., 2004; Lohman and Remington, 2008. Total glutathione concentration was 10mM and a $E_0´$ of -240 mV was considered for the glutathione couple (Bekendam et al., 2016). Fraction of HyPer reduced was determined from the excitation fluorescence intensity ratio 488/405 nm after equilibrium with GSH/GSSG mixtures for 3 h under strictly degassed buffer. Standards fully reduced or fully oxidized were achieved for 10 mM GSH and 10 mM GSSG, respectively.

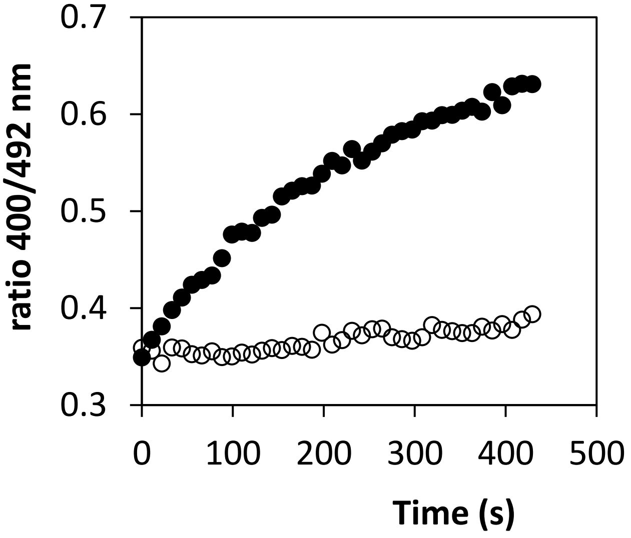


Figure S5. Oxidation of $roGFP2_{RED}$ catalyzed by PDIA1 $WT_{OX}$. roGFP2 was reduced in 50 mM DTT and gel filtered to remove DTT before being mixed with 5 µM PDIA1 $WT_{OX}$ (●) or with buffer (○).

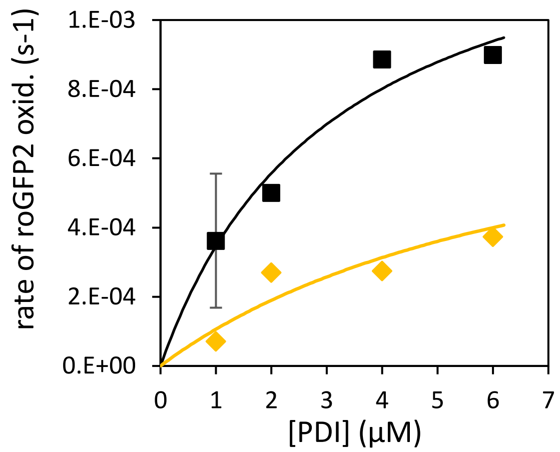


Figure S6. Rate of $roGFP2_{RED}$ oxidation by increasing concentrations of PDIA1 (WT ■; W396A ◆) plus 3 μM ERO1α. PDIA1 was added reduced to prevent roGFP2 oxidation unless PDIA1 is oxidized by ERO1α.

Table S1. Primers to mutagenise PDIA1 showing the mutagenized codons in bold.

| **PDIA1 mutant** | **Primers** |
|---|---|
| C53S | Fw: 5´-TTCTATGCCCCTTGG**TCC**GGCCACTGCAAGGCT-3’<br>Rv: 5´-AGCCTTGCAGTGGCC**GGA**CCAAGGGGCATAGAA-3´ |
| C56S | Fw: 5´-GCCCCTTGGTGTGGCCAC**TCC**AAGGCTCTGGCCCCTGAG-3’<br>Rv: 5´-CTCAGGGGCCAGAGCCTT**GGA**GTGGCCACACCAAGGGGC-3´ |
| C397S | Fw: 5´-GAGTTCTATGCCCCATGG**TCC**GGTCACTGCAAACAGTTG-3’<br>Rv: 5´-CAACTGTTTGCAGTGACC**GGA**CCATGGGGCATAGAACTC-3´ |
| C400S | Fw: 5´-GCCCCATGGTGTGGTCAC**TCC**AAACAGTTGGCTCCCATT-3’<br>Rv: 5´-AATGGGAGCCAACTGTTT**GGA**GTGACCACACCATGGGGC-3´ |
| *a* AS (C397S-C400S) | Fw: 5´-GAGTTCTATGCCCCATGG**TCC**GGTCAC**TCC**AAACAGTTG-3’<br>Rv: 5´-CAACTGTTT**GGA**GTGACC**GGA**CCATGGGGCATAGAACTC-3´ |
| *a´* AS (C53S-C56S) | Fw: 5´-TTCTATGCCCCTTGG**TCC**GGCCAC**TCC**AAGGCTC-3’<br>Rv: 5´-GAGCCTT**GGA**GTGGCC**GGA**CCAAGGGGCATAGAA-3´ |
| W396A | Fw: 5´-CTTTGTGGAGTTCTATGCCCCA**GCG**TGTGGTCACTGCAAACAGTTGG-3’<br>Rv: 5´-CCAACTGTTTGCAGTGACCACA**CGC**TGGGGCATAGAACTCCACAAAG-3´ |
| R300A | Fw: 5´-GACCACACCGACAACCAG**GCC**ATCCTCGAGTTCTTTGGC-3’<br>Rv: 5´-GCCAAAGAACTCGAGGAT**GGC**CTGGTTGTCGGTGTGGTC-3´ |
